# Predictive modeling of virus inactivation by UV

**DOI:** 10.1101/2020.10.27.355479

**Authors:** Nicole C. Rockey, James B. Henderson, Kaitlyn Chin, Lutgarde Raskin, Krista R. Wigginton

**Affiliations:** Department of Civil & Environmental Engineering, University of Michigan, Ann Arbor, MI; Consulting for Statistics, Computing and Analytics Research, University of Michigan, Ann Arbor, MI

**Author notes:** Corresponding author Mailing address: Department of Civil and Environmental Engineering, 1351 Beal Ave., 181 EWRE, Ann Arbor, MI 48109-2125, USA.

## Abstract

Disinfection strategies are commonly applied to inactivate pathogenic viruses in water, food, air, and on surfaces to prevent the spread of infectious diseases. Determining how quickly viruses are inactivated to mitigate health risks is not always feasible due to biosafety restrictions or difficulties with virus culturability. Therefore, methods that would rapidly predict kinetics of virus inactivation by UV_254_ would be valuable, particularly for emerging and difficult-to-culture viruses. We conducted a rapid systematic literature review to collect high-quality inactivation rate constants for a wide range of viruses. Using these data and basic virus information (e.g., genome sequence attributes), we developed and evaluated four different model classes, including linear and non-linear approaches, to find the top performing prediction model. For both the (+) ssRNA and dsDNA virus types, multiple linear regressions were the top performing model classes. In both cases, the cross-validated root mean squared relative prediction errors were similar to those associated with experimental rate constants. We tested the models by predicting and measuring inactivation rate constants for two viruses that were not identified in our systematic review, including a (+) ssRNA mouse coronavirus and a dsDNA marine bacteriophage; the predicted rate constants were within 7% and 71% of the experimental rate constants, respectively. Finally, we applied our models to predict the UV_254_ rate constants of several viruses for which high-quality UV_254_ inactivation data are not available. Our models will be valuable for predicting inactivation kinetics of emerging or difficult-to-culture viruses.

## Introduction

Viruses can cause diverse and costly illnesses in humans and other animals (1). A variety of approaches have therefore been developed to decontaminate food, water, air, and surfaces that may contain infective viruses (2–7). UV_254_ treatment, in particular, is gaining popularity as an alternative to more traditional chemical disinfection strategies (8–10). Viruses can have highly variable UV_254_ susceptibilities (11, 12). For example, two dsDNA viruses, adenovirus type 40 and bacteriophage T6, are inactivated by UV_254_ at the widely varying rates of ∼ 0.06 cm^2^ mJ^−1^ (13–18) and ∼ 5.4 cm^2^ mJ^−1^ (19), respectively.

Viruses have diverse genome types, including double-stranded RNA (dsRNA), single-stranded RNA (ssRNA), double-stranded DNA (dsDNA), and single-stranded DNA (ssDNA). UV_254_ inactivates by primarily targeting viral genetic material, and the different biochemical structures associated with these viral genome types result in distinct sensitivities to UV_254_ (20). Nucleic acid primary structure, or nucleotide base sequence, also affects UV_254_ genome reactivity – pyrimidine bases, for instance, are about an order of magnitude more reactive with UV_254_ than purine bases (21, 22). Different replication modes among viruses can also impact susceptibility to UV_254_. For example, the reverse transcriptase enzymes involved in generation of retrovirus mRNA may have different fidelities to photochemical modifications in nucleic acid compared to the RNA dependent RNA polymerase enzymes used by other RNA viruses to synthesize mRNA (23). Additional differences in viral infection cycles impact virus sensitivity to UV_254_ (24). dsDNA virus genomes, for example, can undergo nucleic acid repair once inside host cells (24–26). This means that a virus may be inactivated by UV_254_ treatment through base modification, only to be repaired and thus rendered infectious again when such repair mechanisms are available. We note these differences in virus genome type and mode of mRNA generation are utilized in the Baltimore virus classification system (e.g., Group 1: dsDNA viruses, Group IV: (+) ssRNA viruses) (1, 27).

Virus disinfection methods are evaluated by enumerating infective viruses before and after treatment, typically with virus culture systems. Relying on culture-based approaches to evaluate inactivation kinetics is often problematic. Most notably, many human viruses that are spread through the environment are not readily culturable. For highly pathogenic viruses that are culturable, disinfection experiments are complicated by biosafety restrictions. Disinfection experiments with severe acute respiratory syndrome (SARS) coronaviruses (SARS-CoV-1 and SARS-CoV-2), for example, are limited to biosafety level 3 laboratories and work with ebolaviruses require biosafety level 4 facilities. Alternative approaches for determining virus inactivation kinetics would be valuable, especially for difficult-to-culture and emerging viruses. Earlier studies have worked towards a predictive manner of evaluating UV_254_ virus inactivation based on virus attributes (28, 29). Recently developed modeling strategies, an improved understanding of virus UV_254_ inactivation mechanisms, and additional high-quality inactivation data published in recent years provide the necessary tools and information to expand upon these initial predictive approaches.

In this study, we develop models to predict rate constants for virus inactivation with UV_254_ treatment in aqueous suspension using variables that are expected to play a role in inactivation, such as genome sequence composition and genome repair information. We conducted a rapid systematic review to gather high quality virus inactivation data from the literature and used the resulting data set to train and validate the predictive performance of four different models (i.e., multiple linear regression, elastic net regularization, boosted trees, and random forests). The models developed in this research will facilitate rapid evaluation of UV_254_ inactivation rate constants for a broad class of virus types based solely on virus genome sequence and genome repair information.

## Results

### Numerous UV_254_ rate constants are available, but only for a limited subset of viruses

We conducted a rapid systematic review to collect UV_254_ inactivation rate constants and used them for the training and validation of models developed to predict virus inactivation kinetics. Of 2,416 initial studies, 531 underwent full text review, and 103 studies were included in the final data set (SI Appendix, Fig. S1). Only data from studies passing a set of experimental criteria (SI Appendix, Supplementary Text) were included to ensure collection of high-quality rate constants. These studies produced 224 experimental inactivation rate constants for 59 viruses (Figure 1; SI Appendix, Table S1). Viruses of different strains and types were considered unique.

**Figure 1.**
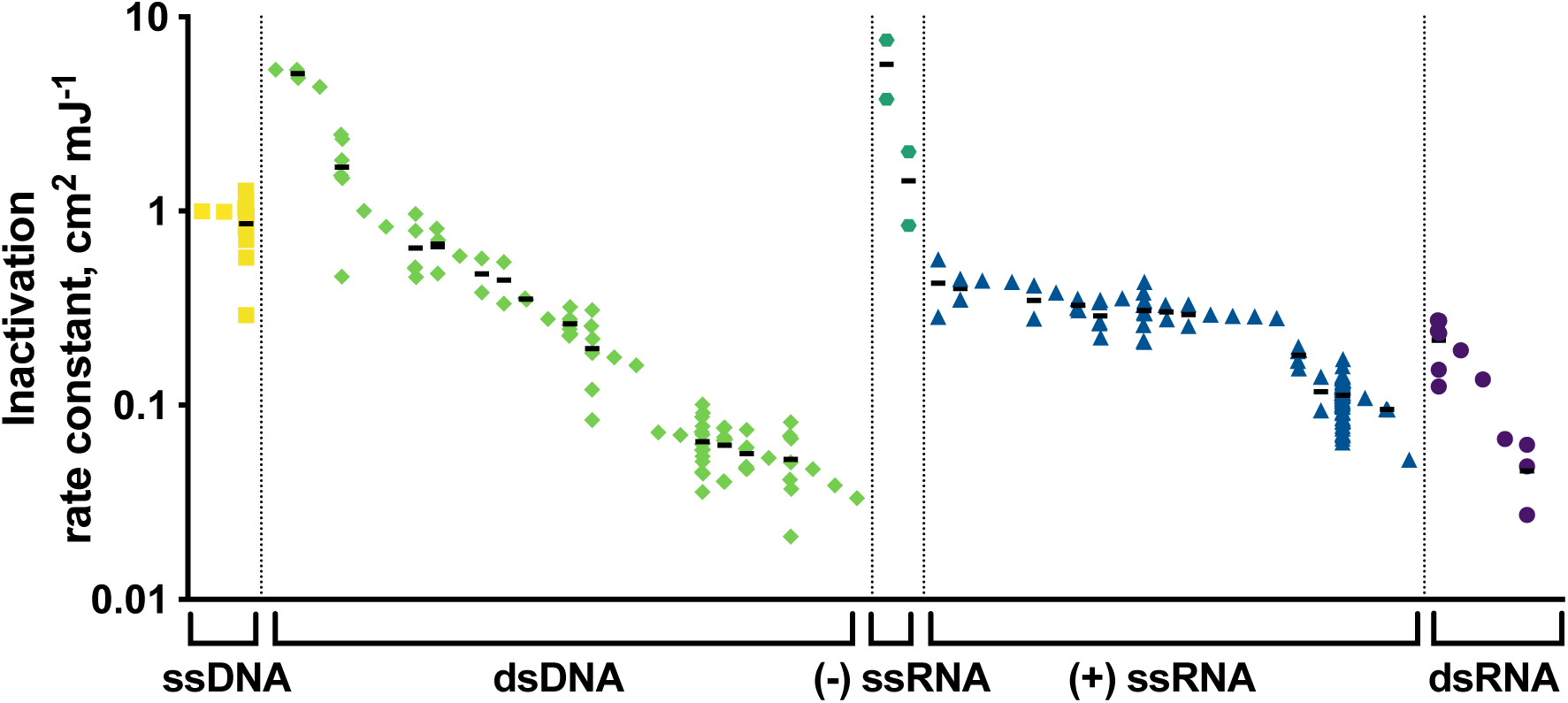
Distribution of UV254 inactivation rate constants collected from the rapid systematic literature review. Black bars denote arithmetic means of inactivation rate constants for viruses with more than one experimental rate constant. Outliers are not included. ssDNA viruses: three viruses, 13 rate constants; dsDNA viruses: 26 viruses,* 84 rate constants; (-) ssRNA viruses: two viruses, four rate constants; (+) ssRNA viruses: 22 viruses, 107 rate constants (four outlier rate constants removed); dsRNA viruses: five viruses, 12 rate constants. Viruses within each Baltimore classification are ordered from highest to lowest mean rate constant from left to right. Rate constants are reported in SI Appendix, Table S1. *Considers two viruses (i.e., adenovirus 5 and adenovirus 41) assayed in host cells with reduced repair abilities as different from the same viruses assayed in wild-type host cells.

More than 350 studies from the full text review that reported conducting UV virus inactivation in aqueous suspension were not included in the final data set. Data were excluded most commonly because the article did not address UV_254_ attenuation in the experimental solution and it could not be ruled out based on details in the materials and methods. Nearly 50% of the extracted rate constants represented only five different viruses. For example, there were 62 different experimental inactivation rates for bacteriophage MS2; in contrast, several viruses, including hepatitis E virus, only had one reported inactivation rate constant, and there were many human viruses with no data that met the review criteria (e.g., influenza viruses, ebolaviruses, coronaviruses, herpesviruses). Ultimately 13, 84, 111, 4, and 12 experimental inactivation rate constants were extracted for ssDNA, dsDNA, (+) ssRNA, (-) ssRNA, and dsRNA viruses, respectively, representing 3, 26, 22, 2, and 5 unique viruses (Figure 1). No rate constants met the inclusion criteria for retroviral (+) ssRNA viruses, referred to as RT-ssRNA viruses. The inactivation rate constants spanned ∼2.5 orders of magnitude (Figure 1) and ranged from 0.021 to 7.6 cm^2^ mJ^−1^. The (-) ssRNA viruses had the largest rate constants on average (k = 3.6 cm^2^ mJ^−1^), while dsRNA viruses had the lowest average rate constants (k = 0.15 cm^2^ mJ^−1^). dsDNA virus constants exhibited the widest range of rate constants, spanning from 0.021 to 5.4 cm^2^ mJ^−1^ with a mean of 0.55 cm^2^ mJ^−1^.

Individual models were developed for the (+) ssRNA and dsDNA virus classes. The limited data sets for viruses in the other Baltimore classifications made it infeasible to develop individual predictive models for the other groups. The data sets used for (+) ssRNA and dsDNA model training and validation included 19 (+) ssRNA viruses with 93 experimental inactivation rate constants and 16 dsDNA viruses with 50 inactivation rate constants, respectively (SI Appendix, Table S1). The model developed with all viruses from the systematic review included 43 viruses with 168 experimental inactivation rate constants.

### Rate constants predicted using common modeling approaches

We used the data collected in the rapid systematic literature review to develop linear regression, elastic net regularization, random forests, and boosted trees models for predicting inactivation rate constants based on several predictors (SI Appendix, Table S2). These model classes were selected to cover a range of different linear and non-linear approaches that are commonly applied in the predictive modeling field (30).

### (+) ssRNA virus model

The cross-validated root mean squared relative prediction errors (RMSrPEs) for the four optimized models varied from 0.22 to 0.95 (Figure 2 and SI Appendix, Table S3), with the top performing multiple linear regression resulting in the lowest RMSrPE out of the four optimized model classes. Various subsets of genomic variables were included in multiple linear regression development. Because these genomic variables are highly collinear, we used principal components that incorporated various genomic variable subsets as predictors in the regression models. Ultimately, the multiple linear regression model with one principal component that incorporated the numbers of cytosines (Cs), uracils (Us), uracil doublets (UUs), and uracil triplets (UUUs) resulted in the lowest RMSrPE (0.22 ± 0.23; RMSrPE ± standard error; SI Appendix, Table S3). Other multiple linear regressions performed similarly (SI Appendix, Table S4). The optimized elastic net regularization and boosted trees models resulted in slightly higher RMSrPEs than the top performing multiple linear regression model (RMSrPE_elastic net_ = 0.28 ± 0.26, RMSrPE_boosted trees_ = 0.32 ± 0.28; SI Appendix, Table S3), and the random forests model had the largest RMSrPE of the (+) ssRNA virus models (RMSrPE_random forests_ = 0.95 ± 0.48; SI Appendix, Table S3). Model performance was significantly reduced in the elastic net and random forests models as compared to the multiple linear regression model (SI Appendix, Table S5).

**Figure 2.**
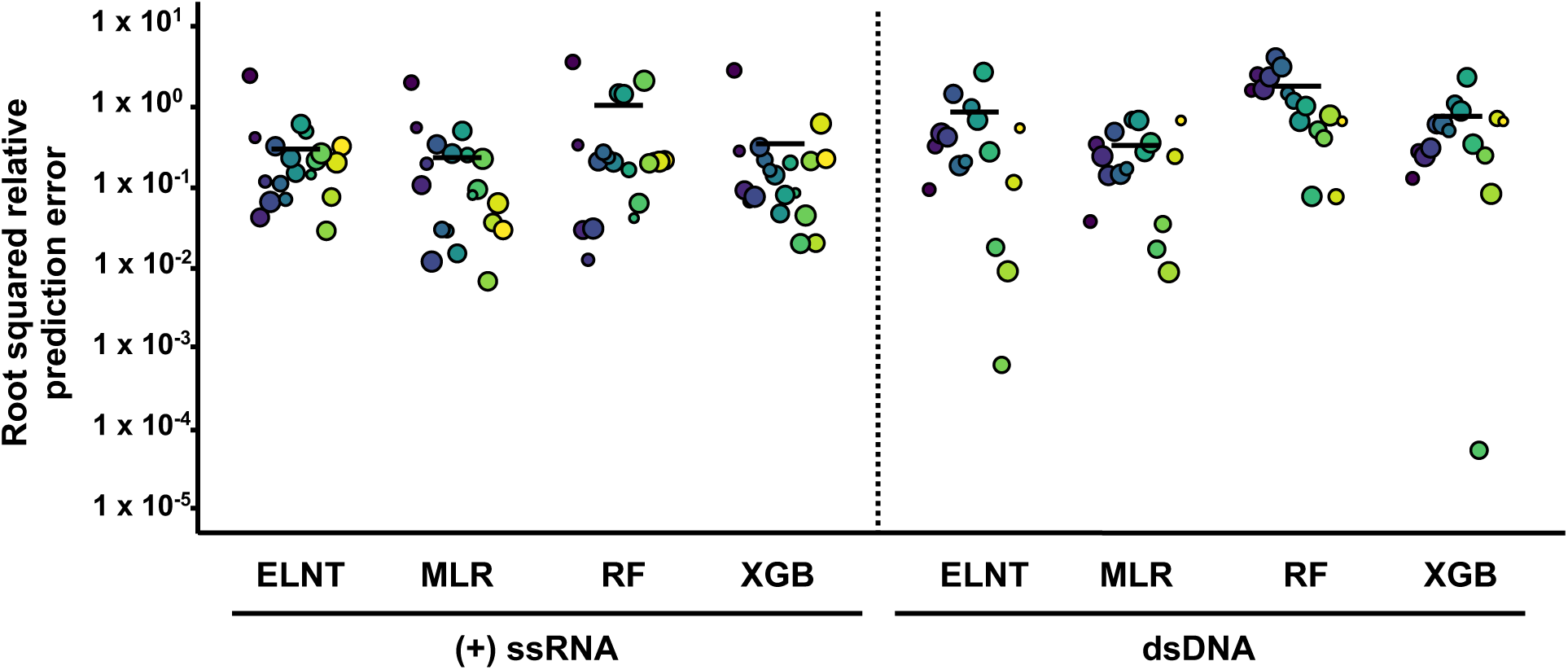
Root squared relative prediction error of virus inactivation rate constants using top performing models from each model class developed with only (+) ssRNA viruses (left) or dsDNA viruses (right) in the training and validation set. Individual symbols indicate the root squared relative prediction error of each virus, and the black bar indicates the model’s root mean squared relative prediction error. Distinct colors represent different viruses, and the symbol sizes represent the weight of the experimental inactivation rate constant used for inverse variance weighting, where a larger symbol indicates a greater weight. MLR = multiple linear regression, ELNT = elastic net regularization, XGB = boosted trees, RF = random forests.

Predicted (+) ssRNA virus rate constants from the top performing model were within 51% of the mean experimental virus inactivation rate constants obtained from the systematic review, with the exception of the rate constant for Atlantic Halibut Nodavirus (percent error = 182%; SI Appendix, Fig. S2a). The RMSrPE from the top performing linear regression model was lower than the estimated relative inter-experimental error of viruses with multiple rate constants in the literature (RMSrPE = 0.22 ± 0.23; relative inter-experimental error = 0.33; Figure 3a). In other words, the predicted rate constants for new (+) ssRNA viruses would be at least as accurate as the rate constants determined through experimental studies.

**Figure 3.**
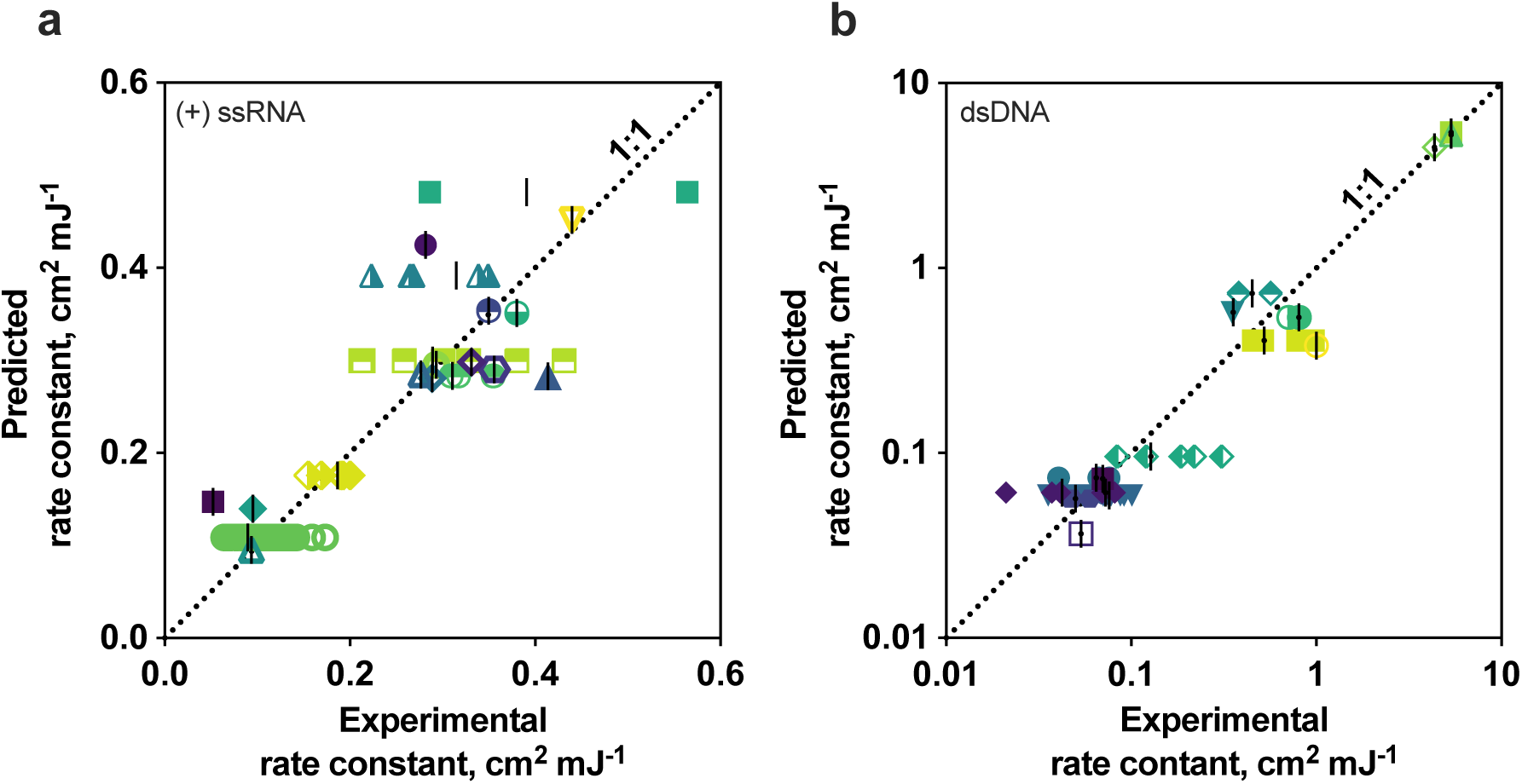
Experimental and predicted cross-validated inactivation rate constants for (+) ssRNA viruses (a) and dsDNA viruses (b) present in the training and validation set. Different colors and symbols represent different viruses. Black lines represent the estimated experimental rate constant for each virus. Data included in the models were obtained from the literature with a rapid systematic review, and all predicted and experimental inactivation rate constants are provided in SI Appendix, Tables S1 and S6.

### dsDNA virus model

The genomic variables used in dsDNA model development were equivalent to the (+) ssRNA models, with the exception that thymines (Ts) were substituted for Us (SI Appendix, Table S1). A major distinction of dsDNA viruses is that their genomes can undergo repair in host cells and this impacts their susceptibility to UV_254_ (24, 31–33). Genome repair can be mediated by the host cell or by viral genes (24), and the varied efficacy of host-mediated dsDNA repair (34–37) impacts virus UV_254_ sensitivity. We included categorical predictors for genome repair mode (i.e., host cell mediated, virus-gene controlled using one repair system, or virus-gene controlled using multiple repair systems) and host cell type (i.e., prokaryotic host, eukaryotic host with wild type repair, or eukaryotic host with reduced repair) in the dsDNA virus inactivation rate constant models. Genome repair mode and host cell type were assigned based on available information and are described in the SI Appendix.

The RMSrPE of the four optimized dsDNA model classes ranged from 0.31 to 1.6 (SI Appendix, Table S3), and the optimized multiple linear regression model outperformed the three other optimized model classes (RMSrPE = 0.31 ± 0.28; Figure 2 and SI Appendix, Table S3). The optimized elastic net and boosted trees RMSrPEs were slightly higher (RMSrPE_elastic net_ = 0.79 ± 0.46, RMSrPE_boosted trees_ = 0.70 ± 0.43), though the difference in model performance was not significant (SI Appendix, Table S5), and the random forests model performed significantly worse (RMSrPE_random forests_ = 1.6 ± 0.66). The top linear regression model included the genome repair mode and host cell type predictors, as well as one principal component comprising the three genomic variables numbers of thymine doublets (TT), thymine quintuplets (TTTTT), and Cs. As with the top-performing (+) ssRNA model, many of the regressions tested with different genomic variable subsets had similar prediction performance, making it difficult to identify which genomic variables were critical for predicting dsDNA virus rate constants (SI Appendix, Table S4). A point estimate comparison of the regression coefficients for the standardized principal component (β_PC1_ = 0.46), genome repair mode (β_genome repair mode_ = 2.7), and host cell type (β_host cell type_ = −0.37) predictors indicates that the genome repair mode predictor is approximately 5.9 times more important than the principal component predictor (β_genome repair mode_/β_PC1_ = 2.7/0.46). Host cell type was comparable in importance to the genomic variable contribution, collectively represented by the principal component. Prediction performance dropped significantly without genome repair mode as a predictor (RMSrPE_opt_ = 0.31 ± 0.28, RMSrPE_no repair_ = 1.0 ± 0.52; SI Appendix, Table S5), further highlighting the importance of genome repair in UV_254_ inactivation.

The multiple linear regression model accurately predicted inactivation rate constants across the wide range of dsDNA virus susceptibilities to UV_254_ (Figure 3b). As with the top performing (+) ssRNA model, the predicted error for the top performing dsDNA model was lower than the estimated inter-experimental error for viruses with more than one experimental rate constant (RMSrPE = 0.31 ± 0.28; inter-experimental error of k_virus_ = 0.45). Predictions were poorest for T7M, B40-8, and lambda predicted (percent error = 62%, 63%, and 62%, respectively; SI Appendix, Fig. S2b), which are bacteriophages with the same form of genome repair mode. The poor prediction of viruses from this group indicates that some of the rate constants in the training data for viruses with these attributes may be inaccurate, leading to worse performance for bacteriophages with host mediated repair.

### All-virus model

Larger data sets generally add predictive power to models, though the increased signal from additional data can be attenuated or negated by increased heterogeneity. We therefore compared the performance of the separate (+) ssRNA and dsDNA virus models with a model that incorporated data from all Baltimore classes. In addition to the genomic variables and repair-related predictors (i.e., genome repair mode and host cell type) included for (+) ssRNA and dsDNA viruses, a categorical predictor for nucleic acid type (i.e., double-stranded or single-stranded) was included. Boosted trees models were the top performing models using all viruses (SI Appendix, Table S3); these performed significantly worse than the models trained using only (+) ssRNA viruses (RMSrPE_(+) ssRNA_ = 0.22 ± 0.23, RMSrPE_all_ = 0.45 ± 0.33; SI Appendix, Table S5) or only dsDNA viruses (RMSrPE_dsDNA_ = 0.31 ± 0.28 vs RMSrPE_all_ = 0.45 ± 0.35; SI Appendix, Tables S3 and S5). This suggests that using our modeling approach and combining viruses with diverse genome types and infection cycles into one model can negatively impact performance of virus predictions, possibly owing to insufficient data from less studied classes. Based on these results, we used the separate (+) ssRNA and dsDNA models for subsequent analyses.

### Predicted rate constants align with new experimental rate constants

We applied the optimized (+) ssRNA and dsDNA models to predict the rate constants of one (+) ssRNA virus and one dsDNA virus for which experimental data were not available and then measured the rate constants experimentally. Specifically, we predicted and measured the rate constants for MHV, a (+) ssRNA mouse coronavirus, and HS2, a dsDNA marine bacteriophage. Based on its large genome size (i.e., ∼ 270% longer than the largest (+) ssRNA virus genome included in the training and validation set) MHV provided an opportunity to assess the (+) ssRNA model’s predictive power using a virus with attributes outside those in the training and validation set (SI Appendix, Fig. S3). HS2 bacteriophage has similar genomic attributes to many of the other viruses in the data set (SI Appendix, Fig. S3), and genome repair-related predictors are the same as those for most of the phages. Bacteriophage MS2 was included in each experimental solution to confirm UV_254_ doses; the measured MS2 rate constants were in line with those in the literature (0.12 to 0.14 cm^2^ mJ^−1^; SI Appendix, Fig. S4 and Table S1).

The predicted inactivation rate constant for MHV (k_pred_ = 2.05 ± 0.88 cm^2^ mJ^−1^; mean ± 95% margin of error) was not significantly different than the experimental rate constant (k_exp_ = 1.92 ± 0.17 cm^−2^ mJ^−1^), with a percent error of only 7% (Figure 4a). The prediction accuracy the model achieved despite MHV’s elevated UV_254_ sensitivity compared with other (+) ssRNA viruses in the data set highlights how linear regression approaches are capable of extrapolating predictions to values distinct from those used in training and validation. In comparison, the MHV inactivation rate constant predicted with the top performing nonlinear approach, boosted trees, was 79% different the experimental value, with a rate constant of 0.40 ± 0.25 cm^2^ mJ^−1^. The accuracy of the MHV rate constant prediction and the relatively low RMSPE obtained for the top performing (+) ssRNA virus model provide confidence that the (+) ssRNA model can effectively predict UV_254_ rate constants for emerging or difficult-to-culture (+) ssRNA viruses.

**Figure 4.**
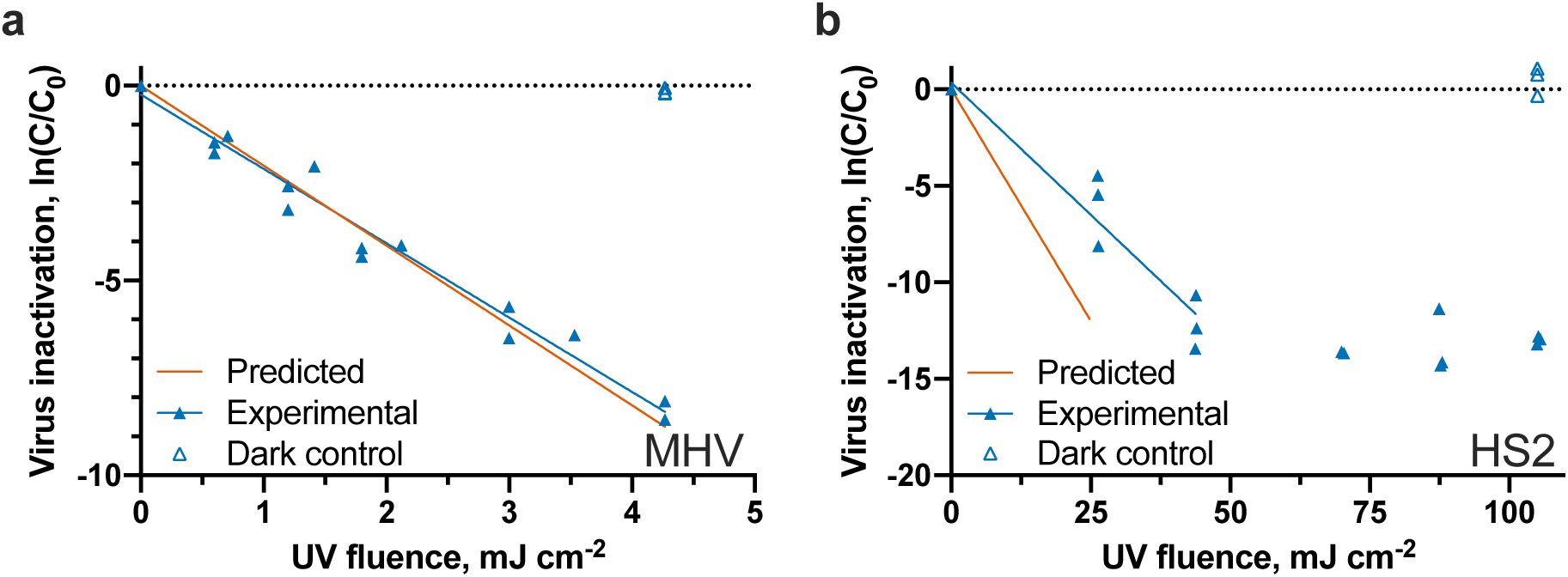
Experimental and predicted UV_254_ inactivation of MHV A59 (a) and HS2 bacteriophage (b). All independent replicates (N = 3) from experiments are shown as individual points. The experimental HS2 inactivation rate constant was determined using the first two UV_254_ fluences due to significant tailing beyond UV_254_ fluences of 50 mJ cm^−2^.

The experimental HS2 inactivation kinetics exhibited significant tailing beyond UV_254_ fluences of 50 mJ cm^−2^; we therefore modeled the first ∼5-log_10_ of inactivation to obtain a rate constant from the first-order portion of the curve. The resulting dsDNA HS2 bacteriophage experimental rate constant of k_exp_ = 0.28 ± 0.08 cm^2^ mJ^−1^ was 71% lower than the predicted rate constant of k_pred_ = 0.48 ± 0.29 cm^2^ mJ^−1^(Figure 4b). Although the error of this dsDNA estimate was larger than that of the (+) ssRNA estimate, the HS2 predicted and experimental constants are not significantly different. This result, in combination with the cross-validation results, suggest that the dsDNA model can effectively predict if a dsDNA virus is particularly resistant to UV_254_ treatment.

### Predictive models estimate inactivation of several emerging and difficult-to-culture viruses

Our systematic review identified a number of important human viruses that lack published high quality UV_254_ inactivation rate constants in the literature. We therefore applied the (+) ssRNA and dsDNA predictive models to estimate the inactivation rates constants for several viruses, including human norovirus, dengue virus, SARS-CoV-2, and several herpesviruses (Table 1). These predictions resulted in a range of inactivation rate constants, from 0.28 for human norovirus to 3.0 cm^2^ mJ^−1^ for human cytomegalovirus.

**Table 1.**
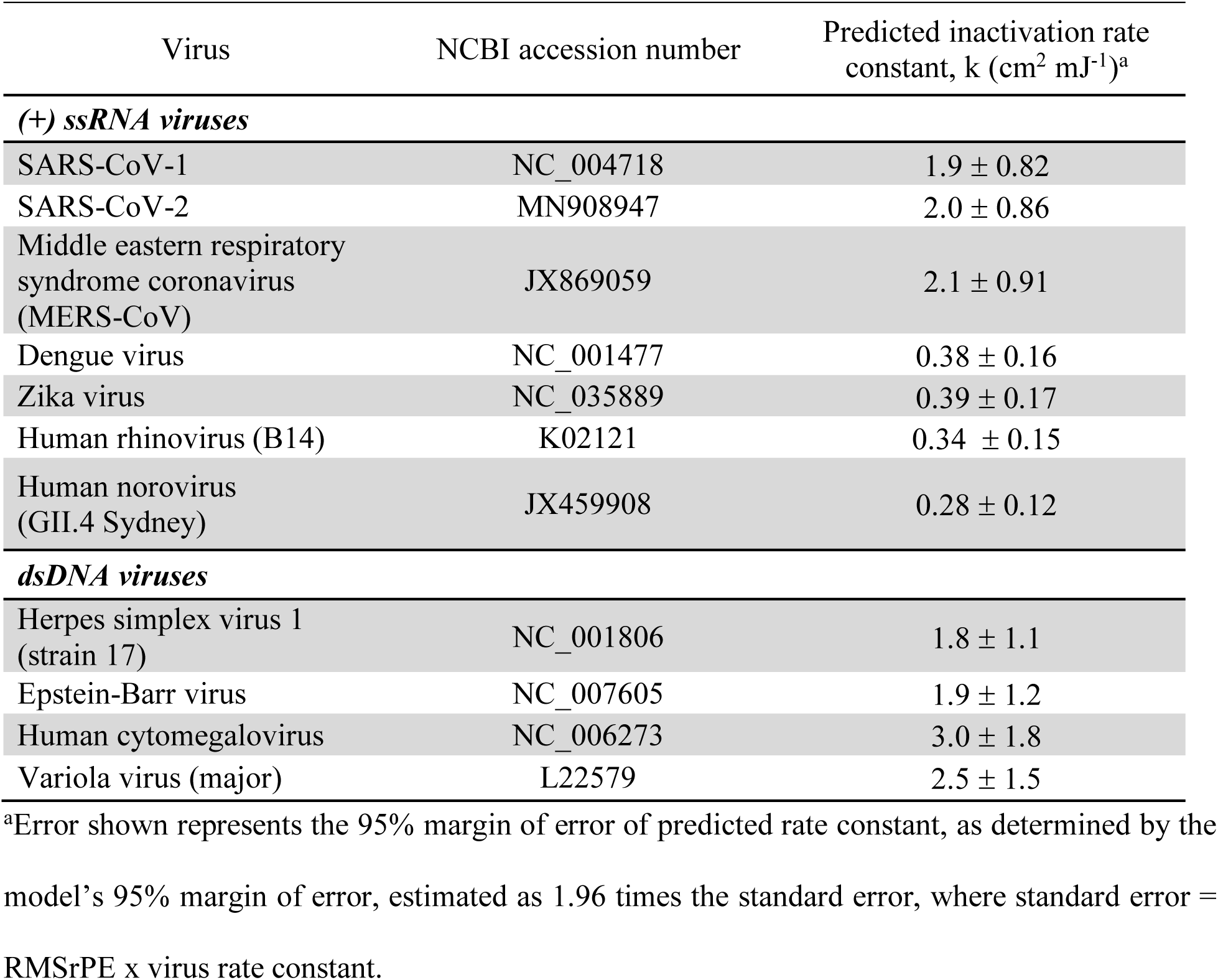
Predicted UV_254_ inactivation rate constants for several viruses without high-quality experimental inactivation rate constants.

| Virus | NCBI accession number | Predicted inactivation rate constant, $k \text{ (cm}^2 \text{ mJ}^{-1})^a$ |
| --- | --- | --- |
| **(+) ssRNA viruses** |  |  |
| SARS-CoV-1 | NC_004718 | $1.9 \pm 0.82$ |
| SARS-CoV-2 | MN908947 | $2.0 \pm 0.86$ |
| Middle eastern respiratory syndrome coronavirus (MERS-CoV) | JX869059 | $2.1 \pm 0.91$ |
| Dengue virus | NC_001477 | $0.38 \pm 0.16$ |
| Zika virus | NC_035889 | $0.39 \pm 0.17$ |
| Human rhinovirus (B14) | K02121 | $0.34 \pm 0.15$ |
| Human norovirus (GII.4 Sydney) | JX459908 | $0.28 \pm 0.12$ |
| **dsDNA viruses** |  |  |
| Herpes simplex virus 1 (strain 17) | NC_001806 | $1.8 \pm 1.1$ |
| Epstein-Barr virus | NC_007605 | $1.9 \pm 1.2$ |
| Human cytomegalovirus | NC_006273 | $3.0 \pm 1.8$ |
| Variola virus (major) | L22579 | $2.5 \pm 1.5$ |

<sup>a</sup>Error shown represents the 95% margin of error of predicted rate constant, as determined by the model's 95% margin of error, estimated as 1.96 times the standard error, where standard error = RMSrPE x virus rate constant.

## Discussion

Through evaluation of a large set of models from four distinct model classes developed with the best currently available data, we identified effective models for predicting UV_254_ inactivation rate constants of (+) ssRNA and dsDNA viruses using simple virus attributes as model predictors. UV_254_ primarily targets viral nucleic acid during irradiation. Pyrimidine bases are more photoreactive than purines (38), and pyrimidine dimers, in particular, cause a large portion of the UV-induced damage to DNA (38–44). Limited research centered on ssRNA photolysis suggests pyrimidine hydrates are the primary lesions inducing UV damage (45). Photochemical damage to nucleic acids can stall or inhibit enzymes required for productive viral infection of host cells (46– 48). Based on this *a priori* knowledge, we included several combinations of pyrimidine bases as predictors in our (+) ssRNA and dsDNA models, namely the numbers of U, UU, UUU, UUUU, UUUUU, C, UC, and CU in (+) ssRNA models and the numbers of T, TT, TTT, TTTT, TTTTT, C, TC, and CT in dsDNA models.

Ultimately, the top performing (+) ssRNA virus model employed one principal component incorporating multiple genomic variables (i.e., numbers of C, U, UU, and UUU), and the top performing dsDNA virus model employed repair mode, host cell type, and one principal component representing three genomic variables (i.e., numbers of C, TT, TTTTT). The relative importance of variables in our top performing predictive models may provide insight into the mechanisms driving UV_254_ inactivation of viruses. Among the (+) ssRNA models, many of the multiple linear regression models that included distinct subsets of genomic variables performed similarly. This is likely because these genomic variables are so highly correlated that different variable combinations resulted in a similar set of principal components as predictors in modeling, ultimately yielding similar performance among different models. Separating the effects of individual genomic variables was therefore difficult in the (+) ssRNA model. Although the top performing model incorporated multiple genomic variables, several linear regression models using as few as one genomic variable as a predictor resulted in similar model performance. This finding demonstrates that simple aspects of the (+) ssRNA genome provide all the necessary information to accurately predict rate constants for this class of viruses. In the dsDNA model, performance was significantly improved when genome repair predictors were included in addition to principal components incorporating genomic variables. The importance of genome repair was expected. For example, the two dsDNA bacteriophages T2 and T4 have similar genome sizes and composition (SI Appendix, Fig. S3b and Table S2) but dissimilar UV_254_ inactivation rate constants (5.1 cm^−2^ mJ^−1^ for T2 and 1.7 cm^−2^ mJ^−1^ for T4; SI Appendix, Table S1). T4 phage’s UV_254_ resistance is due to an additional virus-controlled repair gene in the T4 genome not present in the T2 genome (50, 51). Interestingly, the relative contribution of genomic variables in the dsDNA model was significantly less than the genome repair predictors, which suggests that genome repair is a more important factor in dsDNA UV_254_ inactivation than genomic variables.

Including genome repair as a model predictor presented some limitations. First, the mode and extent of genome repair is not known for many viruses and has not been well-studied across virus families. A single predictor encompassing the contribution of genome repair was therefore not possible. We instead applied multiple categorical predictors. With this approach, only viruses that shared a particular genome repair mode or host cell type with at least one other virus in the dsDNA data set could be used in cross-validation. Ultimately, the data set used for dsDNA model development and validation lacked numerous forms of dsDNA viruses with distinct repair modes and host cell types, resulting in uncertainty in model performance for certain dsDNA viruses not represented in the training and validation set. To improve future dsDNA virus models, it is critical to have a better understanding of genome repair mechanisms and how they affect UV_254_ inactivation.

Our top performing UV_254_ virus prediction models provide improvements over earlier prediction approaches (28, 29). On average, the (+) ssRNA and dsDNA virus models predicted rate constants to within ∼0.2x and ∼0.3x of experimental constants, respectively. A previous approach using genome length to determine genome size-normalized sensitivity values for a number of virus families expected uncertainties in predicted values of ∼2x (28). A more recent approach developed predictive models for ssRNA and dsDNA UV_254_ inactivation using genome dimer formation potential, a value that incorporated pyrimidine doublets, genome length, and purines with adjacent pyrimidine doublets (29). Their reported error as a coefficient of determination (i.e., R^2^) was 0.67 for ssRNA viruses compared to 0.74 (adjusted R^2^) for our model, and an R^2^ value of 0.62 for dsDNA viruses compared to 0.99 (adjusted R^2^) for our model. Several factors can be attributed to the improved performance of our models, including extensive curation of data based on quality and the incorporation of genome repair into dsDNA modeling.

In light of the coronavirus disease 2019 (COVID-19) pandemic and the need for effective decontamination strategies, our predictive models provided an opportunity to predict rate constants for a critical group of viruses with very little published inactivation data. Limited data on UV_254_ inactivation for coronaviruses in aqueous suspension are available and the published information did not pass the inclusion criteria of our rapid systematic review (10, 52–54). This paucity of information on the susceptibility of coronaviruses to UV_254_ is of critical importance for developing effective decontamination strategies. Our predicted rate constants for SARS-CoV-1, SARS-CoV-2, and MERS, and our measured rate constant for the mouse coronavirus MHV, suggest that coronaviruses are much more susceptible to UV_254_ inactivation than other (+) ssRNA viruses. A recent estimate of SARS-CoV-2 UV_254_ susceptibility using the previously developed Lytle and Sagripanti approach (28) is ∼ 1.7x greater than our estimate indicates (55). Discrepancies in new experimental coronavirus data still persist, likely stemming from a lack of checks on UV_254_ attenuation of suspensions.

More robust models are possible with larger data sets that consist of more diverse viruses. Unfortunately, a large portion of UV_254_ inactivation data found during the rapid systematic review did not pass our inclusion criteria. The most common reason for excluding data from our systematic review was a failure to report solution UV_254_ attenuation. An earlier study of SARS-CoV-1 inactivation by UV_254_ (54), for example, did not account for UV_254_ attenuation in the experimental DMEM suspension. The reported inactivation rate constant of 0.003 cm^2^ mJ^−1^ was nearly three orders of magnitude lower than our predicted rate constant for SARS-CoV-1 and our measured value for MHV, likely in part due to solution attenuation. We estimate that their rate constant would be closer to 0.35 cm^2^ mJ^−1^ after accounting for solution attenuation. This value more closely aligns with our coronavirus values. Similarly, several studies reported UV_254_ inactivation of viruses in blood products without describing how attenuation was considered in their reported doses (10, 56–58). Although these doses are likely representative for these fluids, they cannot be extrapolated to other matrices. More stringent reporting of UV_254_ experimental conditions (59), including matrix solution transmission at 254 nm, will facilitate future modeling efforts. We note when UV_254_ inactivation rate constants are known for a solution with 100% transmittance (e.g., purified virus in buffer solution), the rate constant can be adjusted to account for a solution with significant attenuation (e.g., blood products) based on the Beer-Lambert law (60).

The developed models allow us to predict the effectiveness of current UV_254_ treatment strategies on viral pathogens that are difficult or impossible to culture. For example, human norovirus, which causes gastrointestinal disease, is a major target of UV_254_ disinfection processes in water treatment and food processing. Our (+) ssRNA virus model predicts an inactivation rate constant of 0.28 cm^2^ mJ^−1^ for human norovirus GII.4, which is similar to our recently reported rate constant of k = 0.27 cm^2^ mJ^−1^ for human norovirus GII.4 Sydney using RT-qPCR data coupled with a full-genome extrapolation approach (61). This finding indicates that current water treatment guidelines for adequate UV_254_ virus inactivation, which are defined to treat adenovirus 41 (62), are more than sufficient to inactivate human norovirus to acceptable levels. In fact, none of the viruses for which we predicted rate constants had UV_254_ resistance greater than viruses in the *Adenoviridae* family.

The limited and unbalanced data set that we obtained from the systematic review and used in modeling efforts created challenges in our modeling work. Of primary concern, we could not take a commonly used approach to evaluating models, in which a portion of data is held back during model development to assess performance. Holding back the typical 10 – 20% of data would correspond to holding back only two to four viruses from the (+) ssRNA or dsDNA classes for testing. This could result in high variance estimates of prediction performance that would also be highly dependent on the viruses withheld during training. We consequently used leave-one-virus-out cross-validation to more efficiently estimate prediction performance on out of sample data. Another limitation of our models is that they were developed and validated for only (+) ssRNA and dsDNA viruses. Although many human viruses are in these two classes, many emerging and noteworthy human viruses belong to other classes. In particular, the (-) ssRNA virus class includes several important human pathogens, such as lassa virus, nipah virus, influenza virus, and ebolavirus. Since only two (-) ssRNA viruses were included in our data set, we were unable to assess whether inactivation rate constants for viruses in this group could be accurately predicted with our (+) ssRNA model. More high quality UV_254_ experimental inactivation data for a broader set of viruses would facilitate the holdout approach for validating models and the development of models for other virus Baltimore classification groups.

This research demonstrates the value of predictive models for estimating virus fate in various settings. Using readily available viral genome data, we developed models to predict UV_254_ inactivation of (+) ssRNA and dsDNA viruses. The benefits of predictive models are underlined by the ongoing COVID-19 pandemic: access to the biosafety level 3 laboratories required to work with SARS-CoV-2 has been limited and, as a result, few experimental inactivation studies have been performed. Our approach can rapidly determine virus susceptibility to UV_254_ using available genomes, but without relying on culture systems that are often unavailable or difficult to access. Other potential applications of our models including identifying outlier UV_254_ data that are published and predicting potential worst-case scenarios for viruses and their susceptibility to UV_254_. Ultimately, we expect that this predictive modeling approach can be applied to estimate inactivation of microorganisms with other disinfectants and in different settings, such as on surfaces or in air.

## Methods

### Rapid systematic review of UV_254_ virus inactivation data

We used a rapid systematic literature review to capture high quality UV_254_ virus inactivation data (63, 64). Data were extracted from studies if they adhered to all of the following criteria: the UV_254_ lamp fluences were measured and reported; sources emitted UV irradiation principally at wavelengths of 253, 253.7, 254, or 255 nm; viruses were irradiated in a liquid suspension; infective viruses were enumerated with quantitative culture-based approaches (e.g., plaque assay); attenuation through the sample solution was taken into account, or negligible UV_254_ attenuation was reported (transmittance > 95%) or could be assumed based on the reported viral stock purification techniques and matrix solution composition; stirring was reported when attenuation was significant (transmittance < 95%); first-order kinetics were reported or could be confirmed with reported data points for at least two UV_254_ fluences; the first-order inactivation rate constant or log-removal dose (e.g., D_99_) was provided or could be determined with data presented in a plot or table. For publications that contained valuable data, but for which not all criteria could be evaluated, corresponding authors were contacted when possible to inquire about the criteria. For studies that reported multiple UV_254_ inactivation experiments for the same virus (e.g., in different solutions, with multiple UV_254_ sources), we combined all data to determine a single inactivation rate constant with linear regression analysis. All data were re-extracted by a second reviewer and discrepancies were addressed. Additional details of our rapid systematic review process are included in the SI Appendix, Supplementary Text.

### Final data set used in modelin

An inactivation rate constant collected in the rapid systematic review was included in the modeling work if the virus’ genome sequence was available through NCBI and if the error associated with the inactivation rate constant was available. Information on NCBI sequence selection is provided in the SI Appendix, Supplementary Text. For viruses with three or more inactivation rate constants obtained from the systematic review, outlier rate constants (i.e., values lying >1.5 times the interquartile range above the third quartile or below the first quartile) were not included in model development. We calculated the inverse variance weighted mean inactivation rate constant for each virus using the following equation:

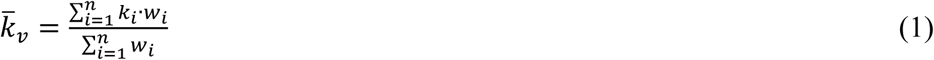

where 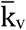 is the inverse variance weighted mean for the virus, n is the number of experimental rate constants for the virus, k_i_ is the inactivation rate constant for experiment i, and w_i_ is the weight for experiment i, defined as:

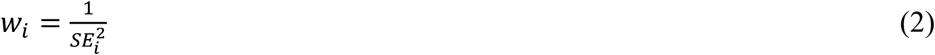

where SE_,i_ is the standard error of the inactivation rate constant for experiment i. The standard error of the inverse variance weighted mean, SE_v_, was evaluated for each virus as:

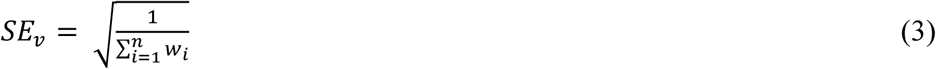

We estimated the inter-experimental error for viruses with more than one experimental rate constant in the literature by determining the residual standard deviation from a weighted least squares regression. Virus was the categorical variable in the regression and experimental rate constant was the dependent variable. Weighting was done using the inverse of the squared experimental standard error normalized by the mean rate constant for that virus.

### Predictors

For model development, we used predictors related to virus structure and behavior that are known or hypothesized to affect UV_254_ inactivation. The specific predictors incorporated included structure of nucleic acid strands (i.e., double-stranded or single stranded), genome length, pyrimidine base content in the genome, sequential pyrimidine bases, genome repair mode, and host cell type. Our reasoning for inclusion of predictors and the methods used to determine values for each predictor are included in the SI. A list of the exact predictors as well as the values used for each virus are available in SI Appendix, Table S2.

### Predictive model optimization

We used four model classes, namely multiple linear regression, elastic net regularization, boosted trees, and random forests, to predict virus inactivation during UV_254_ disinfection. For each model class, we developed individual models using only (+) ssRNA viruses and only dsDNA viruses. We also generated a single model developed using all viruses included in the collected data set and thus not separated by virus Baltimore classification groups. We assessed model performance using leave-one-virus-out cross-validation. Further details of model training, validation, and prediction performance evaluation are included in the SI Appendix, Supplementary Text. Data manipulation, statistical analyses, and modeling work were conducted in R software version 4.0.0 (65). The raw data files and the scripts for model development and prediction will be made available on Github upon publication.

### Multiple linear regression

Several of the genomic variables are collinear (e.g., numbers of U and UU). We therefore conducted principal component analysis (PCA) on the genomic variables prior to linear modeling to reduce variable dimensionality and eliminate collinearity. The predictors nucleic acid type, genome repair mode, and host cell type were not included in the PCA. We then developed linear regression models containing either the first, first and second, or first, second, and third principal components, as well as the other predictors. Only the first through third principal components were assessed for inclusion in the linear regression models, because they cumulatively explained 97% of the variation in genomic variables. Genomic variables were standardized to unit variance prior to PCA to eliminate dissimilarities in the magnitude of variable values. Linear regression can include one or more predictors that can affect model accuracy. We therefore used best subset selection to evaluate a wide range of potential multiple linear regression models.

### Elastic net regularization

As an alternative to best subset selection, we considered linear regression with parameter regularization using L1 (“Lasso”) and L2 (“Ridge”) penalties, a technique known as the elastic net. We used the ‘glmnet’ package in R to create models with elastic net regularization. The alpha and lambda hyperparameters, which control the relative contribution and overall scale of the L1 and L2 penalties, respectively, were tuned using a grid search to find the optimal hyperparameters for the data set as determined by leave-one-virus-out cross-validation. Specifically, 11 different values ranging from 0 to 1 with a step of 0.1 were assessed for the hyperparameter alpha, and 100 different lambda values were evaluated for each alpha.

### Random forests

To accommodate the use of the modified inverse variance weights, the random forests model was developed in R using the ‘xgboost’ package with a single round of boosting, and other hyperparameters were set to match defaults from the ‘randomForest’ package as well as possible (66).

### Boosted trees

Boosted trees modeling was conducted using the ‘xgboost’ package in R. The number of boosting rounds was selected to minimize the cross-validated error. The hyperparameters for learning rate, tree depth, and minimum terminal node weight were 0.3, 6, and 1, respectively.

### Experimental and predicted UV_254_ inactivation of murine hepatitis virus (MHV) and bacteriophage HS2

To consider how well the models may predict inactivation of a virus not already included in the collected data set, we determined the UV_254_ inactivation rate constant of MHV, a virus in the *Coronaviridae* family and *Betacoronavirus* genus, and of HS2, a marine bacteriophage, and compared experimental inactivation to the model’s predicted inactivation. Virus propagation and enumeration details are provided in the SI Appendix, Supplementary Text.

### UV_254_ inactivation of viruses

All UV_254_ inactivation experiments were conducted with a custom-made collimated beam reactor containing 0.16 mW cm^−2^ lamps (model G15T8, Philips). UV_254_ irradiance was determined using chemical actinometry (67, 68) and MS2 (ATCC 15597-B1) was included in all experimental solutions as a biodosimeter to further confirm UV_254_ doses. Infective MS2 was assessed using the double agar overlay approach with host *Escherichia coli* (ATCC 15597) (69). For each UV_254_ exposure, 2 mL of the experimental solution was added to a 10 mL glass beaker and continuously stirred. Sample solution depth (0.8 cm) and transmittance (∼ 47% to 53% for MHV experiments, ∼ 79% to 80% for HS2 experiments) were used to determine the average UV_254_ irradiance of the sample according to the Beer-Lambert law (60). Infective viruses were assayed immediately following experiments. Dark controls were conducted with each experiment and consisted of the virus suspended in experimental solution but stored in the dark on ice for the duration of experiments. Three independent replicates were conducted for each inactivation experiment.

For MHV experiments, solutions contained MHV and MS2 diluted in 1X PBS to a final concentration of ∼ 10^5^ pfu/mL and ∼ 10^10^ pfu/mL, respectively. Samples were exposed to UV_254_ for 0 s, 5 s, 15 s, 25 s, and 35 s, which corresponded to UV_254_ doses of approximately 0 mJ cm^−2^, 0.62 mJ cm^−2^, 1.2 mJ cm^−2^, 1.9 mJ cm^−2^, 3.1 mJ cm^−2^, and 4.3 mJ cm^−2^. MS2 infectivity was assayed after larger UV_254_ doses due to its slower inactivation kinetics, namely 37 mJ cm^−2^ and 74 mJ cm^−2^. For HS2 experiments, solutions contained HS2 and MS2 diluted in 1X PBS to a final concentration of ∼ 10^8^ pfu/mL and ∼ 10^9^ pfu/mL, respectively. Samples were irradiated for 0 s, 180 s, 300 s, 480 s, 600 s, and 720 s, which resulted in UV_254_ doses of approximately 0 mJ cm^−2^, 26 mJ cm^−2^, 44 mJ cm^−2^, 70 mJ cm^−2^, 88 mJ cm^−2^, and 105 mJ cm^−2^.

The inactivation rate constant, k_exp_ in cm^2^ mJ^−1^, for MHV, HS2, and MS2 was determined by the following equation:

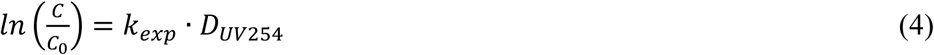

where C_0_ and C are infectious virus concentrations before and after UV_254_ exposure, respectively, in pfu/mL, and D_UV254_ is the average UV_254_ dose, in mJ cm^−2^.

Experimental inactivation rate constants (i.e., k_exp_) were determined with linear regression analyses conducted in Prism version 8.4.2 (GraphPad) to obtain experimental inactivation rate constants (i.e., k_exp_). UV_254_ inactivation curves for some viruses exhibited tailing at high doses. In these situations, only the linear portions of the inactivation curves were included in the linear regression analyses.

### MHV and HS2 inactivation rate constant prediction

The UV_254_ inactivation rate constants of MHV and HS2 were predicted using the best-performing inactivation models for (+) ssRNA viruses and dsDNA viruses, respectively. The MHV genome sequence was provided by Dr. Leibowitz (SI Appendix, Supplementary Text File S1), and the HS2 genome sequence is available in NCBI (accession no. KF302036).

### Predicting UV_254_ inactivation of emerging or difficult-to-culture viruses

The inactivation rates of several emerging and difficult-to-culture viruses, including SARS-CoV-2, were predicted using the best-performing inactivation model. Sequence data for these viruses were obtained from NCBI and all viruses with sequence information are included in SI Appendix, Table S2.

## Acknowledgments

This work was supported by the Water Research Foundation (WRRF #15-07). Nicole C. Rockey was supported by a Rackham Predoctoral Fellowship and a National Science Foundation Graduate Research Fellowship (award no. 2015205675). Kaitlyn Chin was supported by National Science Foundation project #1545756.

## Competing Interests

The authors declare no competing financial interests.

## Notes

### Competing Interest Statement

The authors have declared no competing interest.

## References

1. S. J. Flint, V. R. Racaniello, G. Rall, A. M. Skalka, L. W. Enquist, “Foundations” in Principles of Virology, Volume 1: Molecular Biology, 4th Ed., (ASM Press, 2015), pp. 2–23.

2. K. A. Hirneisen, et al., Viral Inactivation in Foods: A Review of Traditional and Novel Food-Processing Technologies. Compr. Rev. Food Sci. Food Saf. 9, 3–20 (2010).

3. W. A. Rutala, D. J. Weber, Healthcare Infection Control Practices Adviosry Committee, “Guideline for Disinfection and Sterilization in Healthcare Facilities” (2008).

4. A. Adhikari, S. Clark, “Disinfection of Microbial Aerosols” in Modeling the Transmission and Prevention of Infectious Disease, C. J. Hurst, Ed. (Springer International Publishing, 2017), pp. 55–71.

5. M. M. Benjamin, D. F. Lawler, Water quality engineering (Wiley, 2013).

6. Metcalf, et al., Wastewater engineering: treatment and resource recovery, 5th Ed. (McGraw-Hill Education, 2014).

7. D. M. Gunter-Ward, et al., Efficacy of ultraviolet (UV-C) light in reducing foodborne pathogens and model viruses in skim milk. J. Food Process. Preserv. 42, e13485 (2018).

8. M. Pavia, E. Simpser, M. Becker, W. K. Mainquist, K. A. Velez, The effect of ultraviolet-C technology on viral infection incidence in a pediatric long-term care facility. Am. J. Infect. Control 46, 720–722 (2018).

9. D. M. Ward, et al., UV-C treatment on the safety of skim milk: Effect on microbial inactivation and cytotoxicity evaluation. J. Food Process Eng. 42, e12944 (2019).

10. M. Eickmann, et al., Inactivation of Ebola virus and Middle East respiratory syndrome coronavirus in platelet concentrates and plasma by ultraviolet C light and methylene blue plus visible light, respectively. Transfusion 58, 2202–2207 (2018).

11. W. A. M. Hijnen, E. F. Beerendonk, G. J. Medema, Inactivation credit of UV radiation for viruses, bacteria and protozoan (oo)cysts in water: A review. Water Res. 40, 3–22 (2006).

12. A. Shimizu, et al., Human T-cell leukaemia virus type I is highly sensitive to UV-C light. J. Gen. Virol. 85, 2397–2406 (2004).

13. J. G. Jacangelo, P. Loughran, B. Petrik, D. Simpson, C. McIlroy, Removal of enteric viruses and selected microbial indicators by UV irradiation of secondary effluent. Water Sci. Technol. 47, 193–198 (2003).

14. K. G. Linden, J. Thurston, R. Schaefer, J. P. Malley, Enhanced UV Inactivation of Adenoviruses under Polychromatic UV Lamps. Appl. Environ. Microbiol. 73, 7571 LP –7574 (2007).

15. H. Guo, X. Chu, J. Hu, Effect of Host Cells on Low- and Medium-Pressure UV Inactivation of Adenoviruses. Appl. Environ. Microbiol. 76, 7068 LP –7075 (2010).

16. J. A. Thurston-Enriquez, C. N. Haas, J. Jacangelo, K. Riley, C. P. Gerba, Inactivation of Feline Calicivirus and Adenovirus Type 40 by UV Radiation. Appl. Environ. Microbiol. 69, 577–582 (2003).

17. Q. S. Meng, C. P. Gerba, Comparative inactivation of enteric adenoviruses, poliovirus and coliphages by ultraviolet irradiation. Water Res. 30, 2665–2668 (1996).

18. J. Malley, et al., Inactivation of Pathogens with Innovative UV Technologies (American Water Works Association Research Foundation, 2004).

19. S. E. Luria, R. Dulbecco, Genetic Recombinations Leading to Production of Active Bacteriophage from Ultraviolet Inactivated Bacteriophage Particles. Genetics 34, 93–125 (1949).

20. Z. Qiao, Y. Ye, P. H. Chang, D. Thirunarayanan, K. R. Wigginton, Nucleic Acid Photolysis by UV254 and the Impact of Virus Encapsidation. Environ. Sci. Technol. (2018) https://doi.org/10.1021/acs.est.8b02308.

21. K. C. Smith, P. C. Hanawalt, “Photochemistry of the Nucleic Acids” in Molecular Photobiology: Inactivation and Recovery, 1st Ed., B. Horecker, N. O. Kaplan, J. Marmur, Eds. (Academic Press, 1969), pp. 57–84.

22. M. Pearson, H. E. Johns, Suppression of hydrate and dimer formation in ultraviolet-irradiated poly (A + U) relative to poly U. J. Mol. Biol. 20, 215–229 (1966).

23. E. E. Henderson, G. Tudor, J.-Y. Yang, Inactivation of the Human Immunodeficiency Virus Type 1 (HIV-1) by Ultraviolet and X Irradiation. Radiat. Res. 131, 169–176 (1992).

24. W. Harm, Biological effects of ultraviolet radiation (Cambridge University Press, 1980).

25. W. Harm, Gene-controlled reactivation of ultraviolet-inactivated bacteriophage. J. Cell. Comp. Physiol. 58, 69–77 (1961).

26. R. S. Day III, Cellular reactivation of ultraviolet-irradiated human adenovirus 2 in normal and xeroderma pigmentosum fibroblasts. Photochem. Photobiol. 19, 9–13 (1974).

27. D. Baltimore, Expression of animal virus genomes. Bacteriol. Rev. 35, 235–241 (1971).

28. C. D. Lytle, J.-L. Sagripanti, Predicted inactivation of viruses of relevance to biodefense by solar radiation. J. Virol. 79, 14244–14252 (2005).

29. W. J. Kowalski, W. P. Bahnfleth, M. T. Hernandez, A Genomic Model for Predicting the Ultraviolet Susceptibility of Viruses. IUVA News 11, 15–28 (2009).

30. M. Kuhn, K. Johnson, Applied Predictive Modeling (2013) https://doi.org/10.1007/978-1-4614-6849-3.

31. W. Harm, On the relationship between host-cell reactivation and UV-reactivation in UV-inactivated phages. Z. Vererbungsl. 94, 67–79 (1963).

32. C. S. Rupert, W. Harm, “Reactivation After Photobiological Damage” in Advances in Radiation Biology, L. G. Augenstein, R. Mason, M. A. X. B. T.-A. in R. B. Zelle, Eds. (Elsevier, 1966), pp. 1–81.

33. R. S. Day, Studies on Repair of Adenovirus 2 by Human Fibroblasts Using Normal, Xeroderma Pigmentosum, and Xeroderma Pigmentosum Heterozygous Strains. Cancer Res. 34, 1965 LP –1970 (1974).

34. A. J. Rainbow, Defective repair of UV-damaged DNA in human tumor and SV40-transformed human cells but not in adenovirus-transformed human cells. Carcinogenesis 10, 1073–1077 (1989).

35. S. L. MacRae, et al., DNA repair in species with extreme lifespan differences. Aging (Albany. NY). 7, 1171–1184 (2015).

36. E. E. Henderson, Host Cell Reactivation of Epstein-Barr Virus in Normal and Repairdefective Leukocytes. Cancer Res. 38, 3256 LP –3263 (1978).

37. C. D. Lytle, S. A. Aaronson, E. Harvey, Host-cell Reactivation in Mammalian Cells. Int. J. Radiat. Biol. Relat. Stud. Physics, Chem. Med. 22, 159–165 (1972).

38. K. C. Smith, Physical and Chemical Changes Induced in Nucleic Acids by Ultraviolet Light. Radiat. Res. Suppl. 6, 54–79 (1966).

39. R. B. Setlow, W. L. Carrier, The disappearance of thymine dimers from DNA: an error-correcting mechanism. Proc. Natl. Acad. Sci. U. S. A. 51, 226–231 (1964).

40. M. M. Becker, Z. Wang, Origin of ultraviolet damage in DNA. J. Mol. Biol. 210, 429–438 (1989).

41. L. M. Kundu, U. Linne, M. Marahiel, T. Carell, RNA Is More UV Resistant than DNA: The Formation of UV-Induced DNA Lesions is Strongly Sequence and Conformation Dependent. Chem. – A Eur. J. 10, 5697–5705 (2004).

42. W. J. Schreier, et al., Thymine Dimerization in DNA Is an Ultrafast Photoreaction. Science (80-.). 315, 625 LP –629 (2007).

43. M. L. Meistrich, Contribution of thymine dimers to the ultraviolet light inactivation of mutants of bacteriophage T4. J. Mol. Biol. 66, 97–106 (1972).

44. Y. K. Law, R. A. Forties, X. Liu, M. G. Poirier, B. Kohler, Sequence-dependent thymine dimer formation and photoreversal rates in double-stranded DNA. Photochem. Photobiol. Sci. 12, 1431–1439 (2013).

45. G. D. Small, M. Tao, M. P. Gordon, Pyrimidine hydrates and dimers in ultraviolet-irradiated tobacco mosaic virus ribonucleic acid. J. Mol. Biol. 38, 75–87 (1968).

46. R. P. Sinha, D.-P. Häder, UV-induced DNA damage and repair: a review. Photochem. Photobiol. Sci. 1, 225–236 (2002).

47. R. P. Eglin, P. Gugerli, P. Wildy, Ultraviolet Irradiation of Herpes Simplex Virus (Type 1): Delayed Transcription and Comparative Sensitivities of Virus Functions. J. Gen. Virol. 49, 23–31 (1980).

48. F. Yuan, et al., Specificity of DNA Lesion Bypass by the Yeast DNA Polymerase η. J. Biol. Chem. 275, 8233–8239 (2000).

49. M. D. Rosenberg, B. J. Casey, A. J. Holmes, Prediction complements explanation in understanding the developing brain. Nat. Commun. 9, 589 (2018).

50. G. Streisinger, The genetic control of ultraviolet sensitivity levels in bacteriophages T2 and T4. Virology 2, 1–12 (1956).

51. W. Harm, Mutants of phage T4 with increased sensitivity to ultraviolet. Virology 19, 66–71 (1963).

52. A. Pratelli, Canine coronavirus inactivation with physical and chemical agents. Vet. J. 177, 71–79 (2008).

53. M. E. R. Darnell, D. R. Taylor, Evaluation of inactivation methods for severe acute respiratory syndrome coronavirus in noncellular blood products. Transfusion 46, 1770–1777 (2006).

54. M. E. R. Darnell, K. Subbarao, S. M. Feinstone, D. R. Taylor, Inactivation of the coronavirus that induces severe acute respiratory syndrome, SARS-CoV. J. Virol. Methods 121, 85–91 (2004).

55. J.-L. Sagripanti, C. D. Lytle, Estimated Inactivation of Coronaviruses by Solar Radiation With Special Reference to COVID-19. Photochem. Photobiol. 96, 731–737 (2020).

56. H. M. Faddy, et al., Inactivation of dengue, chikungunya, and Ross River viruses in platelet concentrates after treatment with ultraviolet C light. Transfusion 56, 1548–1555 (2016).

57. E. Blázquez, et al., Evaluation of the effectiveness of the SurePure Turbulator ultraviolet-C irradiation equipment on inactivation of different enveloped and non-enveloped viruses inoculated in commercially collected liquid animal plasma. PLoS One 14 (2019).

58. H. Mohr, et al., A novel approach to pathogen reduction in platelet concentrates using short-wave ultraviolet light. Transfusion 49, 2612–2624 (2009).

59. J. R. Bolton, K. G. Linden, Standardization of Methods for Fluence (UV Dose) Determination in Bench-Scale UV Experiments. J. Environ. Eng. 129, 209–215 (2003).

60. H. J. Morowitz, Absorption Effects in Volume Irradiation of Microorganisms. Science (80-.). 111, 229–230 (1950).

61. N. Rockey, et al., UV Disinfection of Human Norovirus: Evaluating Infectivity Using a Genome-Wide PCR-Based Approach. Environ. Sci. Technol. 54, 2851–2858 (2020).

62. U.S. Environmental Protection Agency, “National primary drinking water regulations: The long term 2 enhanced surface water treatment rule. EPA-HQ-” (2006).

63. A. C. Tricco, et al., A scoping review of rapid review methods. BMC Med. 13, 224 (2015).

64. R. Ganann, D. Ciliska, H. Thomas, Expediting systematic reviews: methods and implications of rapid reviews. Implement. Sci. 5, 56 (2010).

65. R Core Team, R: A Language and Environment for Statistical Computing (2020).

66. xgboost developers, Random Forests in XGBoost.

67. R. O. Rahn, J. Bolton, M. I. Stefan, The lodide/lodate Actinometer in UV Disinfection: Determination of the Fluence Rate Distribution in UV Reactors. Photochem. Photobiol. 82, 611–615 (2006).

68. R. O. Rahn, Potassium Iodide as a Chemical Actinometer for 254 nm Radiation: Use of lodate as an Electron Scavenger. Photochem. Photobiol. 66, 450–455 (1997).

69. United States Environmental Protection Agency, “Method 1601: Male-specific (F+) and Somatic Coliphage in Water by Two-step Enrichment Procedure” (2001).

