## Supplemental Information for "Predictive modeling of virus inactivation by UV"

**This PDF file includes:**

Supplementary text
Figs. S1 to S4
Tables S1 to S6

SI References

**Other supplementary materials for this manuscript include the following:**

Supplementary text file S1 (MHV A59 genome)

#### Supplementary Text

***Rapid systematic literature review.*** The Web of Science Core Collection database was used to obtain records for the rapid systematic literature review. Specifically, the database was searched by "Topic" (i.e., search for terms in the title, abstract, author keywords, or Keywords Plus) using the following search term: ((UV OR ultraviolet OR UVC) AND (inactivat* OR disinfect* OR degradat*) AND (virus OR viral OR phage OR bacteriophage)) in August 2019. All records, including conference proceedings and peer-reviewed publications, were output for consideration. Any records containing duplicate content (i.e., duplicate publications, conference proceedings with information also published in a peer-reviewed journal) were removed, and only records in the English language were considered. In addition, any review and microbial risk assessment publications were removed; all references of these papers, however, were screened and relevant references were included in the full-text review (SI Appendix, Fig. S1).

During the first round of screening, the title and abstract of each result were evaluated and all records whose details indicated possible inclusion of $UV_{254}$ virus inactivation data were kept. The text of all records passing the initial screening were then reviewed in full. Any records meeting the inclusion criteria were added to the final study set and $UV_{254}$ virus inactivation data were extracted. For publications that did not allow evaluation of all criteria were not explicitly stated, corresponding authors were contacted when possible to confirm whether remaining criteria were met. Screening and full-text review were carried out by one reviewer.

$UV_{254}$ inactivation rate constants (k), in $cm^2$ $mJ^{-1}$, defined as the natural logarithm transformed reduction in infectious virus concentration per $UV_{254}$ dose, were extracted from the publications. This relationship is described by Chick-Watson kinetics:

$$\ln\left(\frac{C}{C_0}\right) = -k \cdot D_{UV_{254}}$$

where $C_0$ and C are the infectious virus concentrations before and after UV exposure, respectively, and $D_{UV254}$ is the $UV_{254}$ dose, in mJ $cm^{-2}$. When available, inactivation plots from the study showing the reduction in infectious virus concentrations for varying UV doses were digitized using DigitizeIt (1). A linear regression of the digitized data, plotted as the natural logarithm transformed reduction in infectious virus concentrations versus $UV_{254}$ dose, was then conducted using Prism version 8.4.2 (GraphPad, La Jolla, CA). The inactivation rate constant was defined as the slope resulting from the linear regression analysis. Only data following first-order kinetics were included in the linear regression analysis. If multiple different inactivation plots with the same virus were conducted in the same study, the extracted data were combined and analyzed in a single linear regression. When no inactivation plot was available or was too blurry to digitize accurately, but the publication reported some form of the inactivation rate constant, this value was extracted and unit conversions were applied as needed. When only the dose required to achieve a specific log-reduction in infectious virus concentrations was reported (e.g., $D_{90}$ or $D_{99}$ values), this value was converted into the inactivation rate constant using Equation 1.

Standard errors were also collected from studies when possible. For digitized data, the standard error of the slope was obtained from the linear regression. When data could not be digitized, but a rate constant and an associated error estimate were reported, the error estimate was extracted and converted to a standard error. Specifically, if a 95% confidence interval for a rate constant was provided, the confidence interval was divided by four to obtain an estimate of the standard error. Data extraction was conducted by one reviewer. To ensure the quality of extracted data, a second reviewer re-extracted all data from the selected studies. Extracted data were then compared among the two reviewers to confirm consistency.

***Genome sequence selection.*** Genome sequences were determined by searching in NCBI for the virus specified in a study. When no details of the exact virus strain or genotype were provided in the methods of a study, the corresponding author was contacted when possible to determine additional details. Ultimately, the NCBI complete genome sequence with a virus description most closely aligned with the virus description from a study was used. We did not include viruses for which we could not find a complete genome sequence as close to or more specific than the species level described in the study. For example, no full-length genome sequence of the bovine calicivirus serotype used by Malley et al. (2) was found on NCBI, so the inactivation rate constant data for this vesivirus work was not included in model development. Environmental virus isolates without available genome information were not used.

***Predictor selection.*** *Virus nucleic acid type.* The form of virus nucleic acid, either double-stranded or single-stranded, was included as a categorical predictor in the combined models because research has shown that there are significant differences in the rate of photoproduct formation with $UV_{254}$ irradiation (3). For RNA in particular, photoproduct formation with $UV_{254}$ exposure appears to be suppressed significantly in double-stranded nucleic acid compared to single-stranded nucleic acid (3, 4).

*Length*. The length of the viral genome has frequently been associated with $UV_{254}$ virus inactivation (5, 6). While length is not directly a factor in $UV_{254}$ inactivation, longer regions of the genome have more possible reaction sites for $UV_{254}$ photoproduct formation, and so longer genomes indirectly lead to higher reaction rates. Length was determined using virus genome sequences available in NCBI databases.

*Virus genome composition.* Certain components of a virus' primary genomic structure are directly impacted by $UV_{254}$ irradiation. Nucleobases in the genome can be altered through direct photolysis, transforming the nucleic acids into photoproducts that may halt or inhibit translation, transcription, or replication of the viral genome and render the virus noninfectious. In particular, pyrimidine bases are about an order of magnitude more reactive with $UV_{254}$ than purine bases (7). The predictors included in modeling that related to nucleobase composition were therefore exclusively focused on pyrimidine content. Pyrimidine dimers and photohydrates are widely considered the most common products resulting from UV irradiation of nucleic acid. Studies indicate pyrimidine dimers cause a large portion of the UV-induced damage to DNA (8–10), and the formation of thymine dimers, in particular, has been extensively studied (7, 11–13). Research focused on RNA photolysis suggests pyrimidine hydrates are the primary lesions inducing UV damage (14), although the presence of pyrimidine dimers in UV-irradiated RNA has been observed (14, 15). Research has also shown hydrate formation in DNA through UV exposure (16). Presence of flanking pyrimidines next to other pyrimidines is suggested to increase reactivity (9, 10, 13, 17). Considering these findings related to nucleic acid photoreactivity, we selected certain predictors related to the number of specific pyrimidine-based sequence combinations in the virus genome. Specifically, the number of uracil bases (U), cytosine bases (C), uracil doublets (UU), uracil triplets (UUU), uracil quadruplets (UUUU), uracil quintuplets (UUUUU), uracil-cytosine pairs (UC), and cytosine-uracil pairs (CU) were included as predictors for RNA viruses. DNA virus predictors included the number of thymine bases (T), cytosine bases (C), thymine doublets (TT), thymine triplets (TTT), thymine quadruplets (TTTT), thymine quintuplets (TTTTT), thymine-cytosine pairs (TC), and cytosine-thymine pairs (CT). We did not incorporate pyrimidine information including the number of CC, CCC, or CCCC in the genome, because past work indicates photoproducts resulting from these nucleobase combinations are not as prevalent as the other base sequences included in the model (18). Additionally, combinations of bases with purines flanking pyrimidines were not included because of the sparsity of data indicating which precise combinations may lead to photoproducts as well as for simplicity in genomic variable combinations.

All nucleic acid composition values were determined using the same genome sequences used to assess viral genome length. For double-stranded genomes, these variables were counted on both the template and complement strands.

*Genome repair mode.* Genome repair is a process by which genomic lesions in nucleic acid can be repaired through enzymatic activity. To our knowledge, dsDNA viruses are the only class of viruses that can undergo dark genome repair following $UV_{254}$ treatment. Genome repair can be host-mediated or virus-gene controlled. Host cell mediated genome repair occurs for many dsDNA viruses. This is because genome repair enzymes in the host cell, meant to repair host dsDNA, can repair viral dsDNA once the virus genome is in the host cell. In contrast, certain viruses encode one or more repair enzymes in their own virus genome (e.g., T-even and T5 phages); in these cases, genome repair is considered to be virus-gene controlled (19, 20). Theoretically, it would be possible that host cell-mediated repair and virus-gene controlled repair occur simultaneously; in T-even phages known to encode genome repair genes, however, studies have shown that the virus destroys host cell repair mechanisms upon viral entry (20). No viruses are known to undergo simultaneous repair by these distinct genome repair modes, and therefore this combined repair mode was not considered in our models. It is also important to note that no genome photoreactivation was considered in modeling, because these processes only occur when samples are exposed to a nonionizing radiation source following the irradiation treatment (20). In all studies included in our data set, none of the samples were subject to photoreactivation steps.

Based on knowledge of genome repair among viruses from the literature, a categorical predictor, namely genome repair mode, was developed for each virus. The four levels for this predictor were: 0 = ‘host cell mediated,’ 1 = ‘virus-gene controlled using one repair system,’ 2 = ‘no repair,’ or 3 = ‘virus-gene controlled using multiple repair systems.’

All viruses outside of the dsDNA virus Baltimore classification were designated as having no repair; to our knowledge, no evidence of genome repair in the (+) ssRNA, (-) ssRNA, dsRNA, or ssDNA virus classes has been observed. Many dsDNA viruses in our collected data set are known to undergo host cell mediated genome repair, including members of the *Adenoviridae* family, polyomavirus, and lambda phage. Unless past research had reported that a dsDNA virus undergoes virus-gene controlled repair, host cell mediated repair was assumed. This is because eukaryotic and prokaryotic hosts have dsDNA genome repair mechanisms, and there is no reason to believe a dsDNA virus genome would not benefit from these repair systems, unless other virus-mediated activities occur; only T-even phages are known to undergo virus-gene controlled genome repair. As a result, only the T-even bacteriophages T2, T5, and T6 were categorized with ‘virus-gene controlled repair using one repair system.’ One virus, T4 bacteriophage, is known to undergo virus-gene controlled repair with multiple repair systems. Beyond repair mode affecting genome repair capabilities, differences among host cell mediated repair do exist, and we evaluated these differences using another categorical predictor, host cell type.

*Host cell type*. Virus diversity results in significant variability in viral hosts. These hosts differ considerably in several respects; in relation to evaluating $UV_{254}$ inactivation of viruses, we are particularly concerned with the host cell’s ability to repair viral genome damage once the $UV_{254}$-damaged virus has entered the host cell. The efficiency of repair depends on the host cell’s ability to repair genomic material. Eukaryotic and prokaryotic cells contain distinct sets of genes for encoding and producing repair enzymes. Among other dissimilarities, the number, function, and regulation of these repair genes differ (21). Even within the same host type, repair capabilities may differ. For example, studies have shown that in human cells from xeroderma pigmentosum patients, virus $UV_{254}$ sensitivity is significantly increased (e.g., > 3x increase in measured rate constants) compared to in wild-type (i.e., normal) host cells for the same inactivated virus; this is because the cells are deficient in one or more of the common repair genes needed to effectively repair dsDNA inside the host (19, 22–25). In addition, work has shown that genome repair systems in the cells of longer-lived mammals (e.g., human) are significantly upregulated compared to those in shorter-lived mammals (i.e., mice) (26). To incorporate these differences in host cell type that impact genome repair, we developed a categorical predictor for host cell type. Three different categories were used, including 0 = 'prokaryotic cells', 1= 'eukaryotic cells with reduced repair', and 2 = 'eukaryotic cells with wild-type repair.'

All bacteriophages were assigned the category for 'prokaryotic cells.' Some experiments with human viruses were assayed in cell lines known to have reduced repair capabilities compared to wild-type human cell lines; these virus experiments were assessed as having 'eukaryotic cells with reduced repair.' The only virus in the modeling data set with this form of repair was human polyomavirus, assayed in SVG-A cells. A previous study showed reduced repair of DNA damage in SV-derived cells but not in other cells evaluated (25). While two adenoviruses in the data set were also evaluated in cell lines with reduced repair (22), these rate constants did not have associated errors and were therefore not included in modeling work. All other dsDNA human viruses were given the host cell type 'eukaryotic cells with wild-type repair.'

**Model training, validation, and prediction performance evaluation.** Inverse relative variance weighting was used in model training, validation and prediction performance evaluation. The weight for each virus, $w_v$, was determined as follows:

$$w_v = \frac{\frac{\bar{k}_v^{\,2}}{SE_v^2}}{\sum_{v=1}^{n_v} \frac{\bar{k}_v^{\,2}}{SE_v^2}}$$

where $n_v$ is the number of viruses in the data set, $\bar{k}_v$ is the inverse variance weighted mean inactivation rate constant for virus v, and $SE_v$ is the standard error of the inverse variance weighted mean inactivation rate constant for virus v.

We employed weighted root mean squared relative prediction error (RMSrPE) to assess model prediction efficacy. Leave-one-virus-out cross-validation was used to determine each model's RMSrPE. Specifically, data from the final curated data set were split into a training set and a validation set for each round of cross-validation so each virus was left out of the training set exactly one time. Weights for viruses in the training set were rescaled to sum to the number of viruses – 1. The squared predicted error was determined for the held-out virus inactivation rate constant in each fold. The resulting relative errors for each virus were weighted by the inverse relative variance weights and the weighted mean computed. The root of these values was then taken to obtain the RMSrPE for a particular model, as shown in the follow equation:

$$RMSrPE = \sqrt{\sum_{v=1}^{n_v} \left[ \left( \bar{k}_v - \hat{k}_v \right)^2 \cdot w_v \right]}$$

where $\hat{k}_v$ is the predicted inactivation rate constant for virus v. The RMSrPE for each model was compared and top performing models were selected based on minimum RMSrPE values. The standard error associated with the relative RMSrPE was determined as the bias corrected weighted sample variance:

$$Standard\ error\ (RMSrPE) = \frac{\sum_{v=1}^{n_v}\left[\left(\bar{k}_v - \hat{k}_v\right)^2 \cdot w_v\right]}{\left(1 - \sum_{v=1}^{n_v} w_v^2\right)}$$

Pairwise comparisons of model performance were conducted using weighted least squares regression to compare the expected log ratio of the squared prediction errors. Specifically, for each virus the squared prediction error determined during cross-validation were transformed to the natural logarithm scale and differenced. A logarithm transform was used to stabilize the variance estimates of large individual prediction errors in some models. A weighted regression was then conducted using only an intercept as the predictor and the transformed squared error ratios as the dependent variable (i.e., a weighted one-sample t-test of the logarithm squared error ratios). The average squared error ratio (ASER) was determined by exponentiating the estimated intercept from this regression. Differences in model performance were considered significant if the exponentiated 95% confidence interval of the ASER did not include 1.

***Virus propagation and enumeration.*** *MHV*. MHV strain A59 was propagated and quantified in delayed brain tumor (DBT) cells (kindly provided by Dr. Julian Leibowitz at Texas A&M Health Science Center College of Medicine) according to published protocols with slight modifications (27, 28). Briefly, DBT cells were grown in medium comprised of Dulbecco's modified Eagle's medium (DMEM) with 4.5 g/L glucose without L-glutamine (Cat. No. 12614F, Lonza), 10% horse serum (Cat. No. 26050088, Life Technologies), 1% penicillin streptomycin (Cat. No. 15140122, Invitrogen), and 1% L-glutamine (100X; Cat. No. 25030081, Invitrogen) at 37°C and 5% $CO_2$. The medium was replaced every 48 to 72 hours.

MHV stocks were generated in 80% confluent DBT cell monolayers at a multiplicity of infection of approximately 0.01. Following 18 to 24 hours of incubation, infected DBT cells were centrifuged at 3,000 x g for 15 minutes at 4ºC and the supernatant was collected. MHV A59 stocks (~$10^6$ pfu/mL) were filter-sterilized with a 0.22 µm sterile polyethersulfone (PES) membrane (Cat. No. 229747, CELLTREAT Scientific) and stored in single-use aliquots at -80ºC.

MHV was enumerated via plaque assay. Specifically, DBT cell monolayers were seeded in 12-well plates (Cat. No. 353043, Corning) and incubated at 37°C and 5% $CO_2$ prior to infection with MHV samples. Plaque assays were performed by inoculating 90% confluent cells with MHV samples diluted in DMEM2 (DMEM with 2% horse serum, 1% penicillin streptomycin, and 1% L-glutamine) for one hour. After inoculation, virus suspensions were removed and replaced with a 1:1 solution of 1.6% agarose (Cat. No. BP160-100, ThermoFisher) and 2xMEM (2x E-MEM (Cat No. 115073101, Quality Biological, Inc.), 5% horse serum, 10 mM HEPES (Cat. No. 17737E, Lonza), 1X MEM non-essential amino acids (Cat. No. 11140050, Invitrogen), 2% L-glutamine, and 2% penicillin streptomycin). Infected DBT cell monolayers were incubated for 48 hours. Plaques were enumerated using neutral red staining (Cat. No. N2889, Sigma-Aldrich) at a final 0.01% concentration in 1X phosphate buffered saline (PBS; Cat. No. 10010023, Invitrogen). Samples were enumerated in triplicate and negative media controls were plated with samples.

*HS2 bacteriophage.* HS2 marine bacteriophage and its host *Pseudoalteromonas* 13-15 (kindly provided by Dr. Melissa Duhaime at the University of Michigan Department of Ecology and Evolutionary Biology) were propagated and enumerated using established methods with modifications (29). Briefly, HS2 bacteriophage stock was generated using the soft agar overlay method. Specifically, soft seawater agar (5 g/L peptone, 1 g/L yeast extract, 10% Widdel salt solution, 0.6% agar) containing HS2 bacteriophage and host bacteria was overlaid on hard seawater agar plates (1 g/L peptone, 0.2 g/L yeast extract, 10% Widdel salt solution, 1.2% agar) and incubated overnight at 25ºC. The soft seawater agar with virus was then scraped off and diluted with SM buffer (100 mM NaCl, 81.2 mM $MgSO_4{\bullet}7H_2O$, 50mM Tris-HCl (pH 7.5)). Chloroform was added to the agar solution (5 mL chloroform per 50 mL solution) and centrifuged at 3000 x g for 10 minutes. The supernatant was aerated to remove residual chloroform and filtered through a 0.45 μm PES membrane. The resulting HS2 bacteriophage stock (~ $10^{11}$ pfu/mL) was stored at 4ºC until use. HS2 infectivity was quantified by plaque assay.

#### References

1. ,  DigitizeIt - Digitizer Software.

2. J. Malley, *et al.*, *Inactivation of Pathogens with Innovative UV Technologies* (American Water Works Association Research Foundation, 2004).

3. M. Pearson, H. E. Johns, Suppression of hydrate and dimer formation in ultraviolet-irradiated poly (A + U) relative to poly U. *J. Mol. Biol.* **20**, 215–229 (1966).

4. Z. Qiao, Y. Ye, P. H. Chang, D. Thirunarayanan, K. R. Wigginton, Nucleic Acid Photolysis by UV254 and the Impact of Virus Encapsidation. *Environ. Sci. Technol.* (2018) https:/doi.org/10.1021/acs.est.8b02308.

5. J. Simonet, C. Gantzer, Inactivation of Poliovirus 1 and F-Specific RNA Phages and Degradation of Their Genomes by UV Irradiation at 254 Nanometers. *Appl. Environ. Microbiol.* **72**, 7671 LP – 7677 (2006).

6. C. D. Lytle, J.-L. Sagripanti, Predicted inactivation of viruses of relevance to biodefense by solar radiation. *J. Virol.* **79**, 14244–14252 (2005).

7. K. C. Smith, Physical and Chemical Changes Induced in Nucleic Acids by Ultraviolet Light. *Radiat. Res. Suppl.* **6**, 54–79 (1966).

8. R. B. Setlow, W. L. Carrier, The disappearance of thymine dimers from DNA: an error-correcting mechanism. *Proc. Natl. Acad. Sci. U. S. A.* **51**, 226–231 (1964).

9. M. M. Becker, Z. Wang, Origin of ultraviolet damage in DNA. *J. Mol. Biol.* **210**, 429–438 (1989).

10. L. M. Kundu, U. Linne, M. Marahiel, T. Carell, RNA Is More UV Resistant than DNA: The Formation of UV-Induced DNA Lesions is Strongly Sequence and Conformation Dependent. *Chem. – A Eur. J.* **10**, 5697–5705 (2004).

11. W. J. Schreier, *et al.*, Thymine Dimerization in DNA Is an Ultrafast Photoreaction. *Science (80-. ).* **315**, 625 LP – 629 (2007).
12. M. L. Meistrich, Contribution of thymine dimers to the ultraviolet light inactivation of mutants of bacteriophage T4. *J. Mol. Biol.* **66**, 97–106 (1972).
13. Y. K. Law, R. A. Forties, X. Liu, M. G. Poirier, B. Kohler, Sequence-dependent thymine dimer formation and photoreversal rates in double-stranded DNA. *Photochem. Photobiol. Sci.* **12**, 1431–1439 (2013).
14. G. D. Small, M. Tao, M. P. Gordon, Pyrimidine hydrates and dimers in ultraviolet-irradiated tobacco mosaic virus ribonucleic acid. *J. Mol. Biol.* **38**, 75–87 (1968).
15. K. B. Freeman, P. V Hariharan, H. E. Johns, The ultraviolet photochemistry of cytidylyl-(3′-5′)-cytidine. *J. Mol. Biol.* **13**, 833-IN16 (1965).
16. G. J. Fisher, H. E. Johns, “4 - Pyrimidine Photohydrates” in S. Y. B. T.-P. and P. of N. A. Wang, Ed. (Academic Press, 1976), pp. 169–224.
17. M. Owada, S. Ihara, K. Toyoshima, Y. Kozai, Y. Sugino, Ultraviolet inactivation of avian sarcoma viruses: Biological and biochemical analysis. *Virology* **69**, 710–718 (1976).
18. T. Douki, J. Cadet, Individual Determination of the Yield of the Main UV-Induced Dimeric Pyrimidine Photoproducts in DNA Suggests a High Mutagenicity of CC Photolesions. *Biochemistry* **40**, 2495–2501 (2001).
19. C. S. Rupert, W. Harm, “Reactivation After Photobiological Damage” in *Advances in Radiation Biology*, L. G. AUGENSTEIN, R. MASON, M. A. X. B. T.-A. in R. B. ZELLE, Eds. (Elsevier, 1966), pp. 1–81.
20. W. Harm, *Biological effects of ultraviolet radiation* (Cambridge University Press, 1980).
21. G. M. Cooper, R. E. Hausman 1947-2015., *The cell : a molecular approach*, 5th ed.

(ASM Press ; Sinauer Associates, 2009).

22. H. Guo, X. Chu, J. Hu, Effect of Host Cells on Low- and Medium-Pressure UV Inactivation of Adenoviruses. *Appl. Environ. Microbiol.* **76**, 7068 LP – 7075 (2010).

23. R. S. Day III, Cellular reactivation of ultraviolet-irradiated human adenovirus 2 in normal and xeroderma pigmentosum fibroblasts. *Photochem. Photobiol.* **19**, 9–13 (1974).

24. A. J. Rainbow, Reduced Capacity to Repair Irradiated Adenovirus in Fibroblasts from Xeroderma Pigmentosum Heterozygotes. *Cancer Res.* **40**, 3945 LP – 3949 (1980).

25. A. J. Rainbow, Defective repair of UV-damaged DNA in human tumor and SV40-transformed human cells but not in adenovirus-transformed human cells. *Carcinogenesis* **10**, 1073–1077 (1989).

26. S. L. MacRae, *et al.*, DNA repair in species with extreme lifespan differences. *Aging (Albany. NY).* **7**, 1171–1184 (2015).

27. J. Leibowitz, G. Kaufman, P. Liu, Coronaviruses: propagation, quantification, storage, and construction of recombinant mouse hepatitis virus. *Curr. Protoc. Microbiol.* **Chapter 15**, Unit-15E.1 (2011).

28. Y. Ye, R. M. Ellenberg, K. E. Graham, K. R. Wigginton, Survivability, Partitioning, and Recovery of Enveloped Viruses in Untreated Municipal Wastewater. *Environ. Sci. Technol.* **50**, 5077–5085 (2016).

29. M. B. Duhaime, *et al.*, Comparative Omics and Trait Analyses of Marine Pseudoalteromonas Phages Advance the Phage OTU Concept. *Front. Microbiol.* **8** (2017).

30. N. Nwachuku, C. P. Gerba, A. Oswald, F. D. Mashadi, Comparative Inactivation of Adenovirus Serotypes by UV Light Disinfection. *Appl. Environ. Microbiol.* **71**, 5633–5636 (2005).

31. N. A. Ballesster, J. P. J. Malley, Sequential disinfection of adenovirus type 2 with UV-chlorine-chloramine. *J. Am. Water Work. Assoc.* **96** (2004).

32. C. S. Baxter, R. Hofmann, M. R. Templeton, M. Brown, R. C. Andrews, Inactivation of Adenovirus Types 2, 5, and 41 in Drinking Water by UV Light, Free Chlorine, and Monochloramine. *J. Environ. Eng.* **133**, 95–103 (2007).

33. S. E. Beck, *et al.*, Wavelength Dependent UV Inactivation and DNA Damage of Adenovirus as Measured by Cell Culture Infectivity and Long Range Quantitative PCR. *Environ. Sci. Technol.* **48**, 591–598 (2014).

34. F. Bosshard, F. Armand, R. Hamelin, T. Kohn, Mechanisms of Human Adenovirus Inactivation by Sunlight and UVC Light as Examined by Quantitative PCR and Quantitative Proteomics. *Appl. Environ. Microbiol.* **79**, 1325 LP – 1332 (2013).

35. S. Bounty, R. A. Rodriguez, K. G. Linden, Inactivation of adenovirus using low-dose UV/H2O2 advanced oxidation. *Water Res.* **46**, 6273–6278 (2012).

36. B. Calgua, *et al.*, UVC Inactivation of dsDNA and ssRNA Viruses in Water: UV Fluences and a qPCR-Based Approach to Evaluate Decay on Viral Infectivity. *Food Environ. Virol.* **6**, 260–268 (2014).

37. A. C. Eischeid, J. N. Meyer, K. G. Linden, UV Disinfection of Adenoviruses: Molecular Indications of DNA Damage Efficiency. *Appl. Environ. Microbiol.* **75**, 23 LP – 28 (2009).

38. C. P. Gerba, D. M. Gramos, N. Nwachuku, Comparative Inactivation of Enteroviruses and Adenovirus 2 by UV Light. *Appl. Environ. Microbiol.* **68**, 5167 LP – 5169 (2002).

39. K. G. Linden, J. Thurston, R. Schaefer, J. P. Malley, Enhanced UV Inactivation of Adenoviruses under Polychromatic UV Lamps. *Appl. Environ. Microbiol.* **73**, 7571 LP – 7574 (2007).

40. R. A. Rodríguez, S. Bounty, K. G. Linden, Long-range quantitative PCR for determining inactivation of adenovirus 2 by ultraviolet light. *J. Appl. Microbiol.* **114**, 1854–1865 (2013).

41. H. Ryu, J. L. Cashdollar, G. S. Fout, K. A. Schrantz, S. Hayes, Applicability of integrated cell culture quantitative PCR (ICC-qPCR) for the detection of infectious adenovirus type 2 in UV disinfection studies. *J. Environ. Sci. Heal. Part A* **50**, 777–787 (2015).

42. G.-A. Shin, K. G. Linden, M. D. Sobsey, Low pressure ultraviolet inactivation of pathogenic enteric viruses and bacteriophages. *J. Environ. Eng. Sci.* **4**, S7–S11 (2005).

43. S. S. Thompson, *et al.*, Detection of Infectious Human Adenoviruses in Tertiary-Treated and Ultraviolet-Disinfected Wastewater. *Water Environ. Res.* **75**, 163–170 (2003).

44. B. Vazquez-Bravo, K. Gonçalves, J. L. Shisler, B. J. Mariñas, Adenovirus Replication Cycle Disruption from Exposure to Polychromatic Ultraviolet Irradiation. *Environ. Sci. Technol.* **52**, 3652–3659 (2018).

45. S. Rattanakul, K. Oguma, H. Sakai, S. Takizawa, Inactivation of Viruses by Combination Processes of UV and Chlorine. *J. Water Environ. Technol.* **12**, 511–523 (2014).

46. S. Rattanakul, K. Oguma, S. Takizawa, Sequential and Simultaneous Applications of UV and Chlorine for Adenovirus Inactivation. *Food Environ. Virol.* **7**, 295–304 (2015).

47. J. Sangsanont, H. Katayama, F. Kurisu, H. Furumai, Capsid-Damaging Effects of UV Irradiation as Measured by Quantitative PCR Coupled with Ethidium Monoazide Treatment. *Food Environ. Virol.* **6**, 269–275 (2014).

48. J. G. Jacangelo, P. Loughran, B. Petrik, D. Simpson, C. McIlroy, Removal of enteric viruses and selected microbial indicators by UV irradiation of secondary effluent. *Water Sci. Technol.* **47**, 193–198 (2003).

49. Q. S. Meng, C. P. Gerba, Comparative inactivation of enteric adenoviruses, poliovirus and coliphages by ultraviolet irradiation. *Water Res.* **30**, 2665–2668 (1996).

50. J. A. Thurston-Enriquez, C. N. Haas, J. Jacangelo, K. Riley, C. P. Gerba, Inactivation of Feline Calicivirus and Adenovirus Type 40 by UV Radiation. *Appl. Environ. Microbiol.* **69**, 577–582 (2003).

51. N. Ding, S. A. Craik, X. Pang, B. Lee, N. F. Neumann, Assessing UV Inactivation of Adenovirus 41 Using Integrated Cell Culture Real-Time qPCR/RT-qPCR. *Water Environ. Res.* **89**, 323–329 (2017).

52. G. Ko, T. L. Cromeans, M. D. Sobsey, UV inactivation of adenovirus type 41 measured by cell culture mRNA RT-PCR. *Water Res.* **39**, 3643–3649 (2005).

53. D. Diston, J. E. Ebdon, H. D. Taylor, The effect of UV-C radiation (254 nm) on candidate microbial source tracking phages infecting a human-specific strain of Bacteroides fragilis (GB-124). *J. Water Health* **10**, 262–270 (2012).

54. R. Sommer, T. Haider, A. Cabaj, W. Pribil, M. Lhotsky, Time dose reciprocity in UV disinfection of water. *Water Sci. Technol.* **38**, 145–150 (1998).

55. R. Sommer, *et al.*, Inactivation of bacteriophages in water by means of non-ionizing (uv-253.7nm) and ionizing (gamma) radiation: a comparative approach. *Water Res.* **35**, 3109–3116 (2001).

56. H. S. Lee, M. D. Sobsey, Survival of prototype strains of somatic coliphage families in environmental waters and when exposed to UV low-pressure monochromatic radiation or heat. *Water Res.* **45**, 3723–3734 (2011).

57. K. D. Martignoni, I. Haselbacher, Inactivation of Bacteriophage Lambda by Combined X-ray and U.V.-light Exposure. *Int. J. Radiat. Biol. Relat. Stud. Physics, Chem. Med.* **35**,

441–447 (1979).

58. A. M. Rauth, The Physical State of Viral Nucleic Acid and the Sensitivity of Viruses to Ultraviolet Light. *Biophys. J.* **5**, 257–273 (1965).

59. R. Latarjet, R. Cramer, L. Montagnier, Inactivation, by UV-, X-, and γ-radiations, of the infecting and transforming capacities of polyoma virus. *Virology* **33**, 104–111 (1967).

60. K. S. Bae, G.-A. Shin, Inactivation of various bacteriophages by different ultraviolet technologies: Development of a reliable virus indicator system for water reuse. *Environ. Eng. Res.* **21**, 350–354 (2016).

61. D. Gerrity, H. Ryu, J. Crittenden, M. Abbaszadegan, Photocatalytic inactivation of viruses using titanium dioxide nanoparticles and low-pressure UV light. *J. Environ. Sci. Heal. Part A* **43**, 1261–1270 (2008).

62. R. D. Kashinkunti, *et al.*, Investigating Multibarrier Inactivation for Cincinnati—UV, By-Products, and Biostability. *J. - AWWA* **96**, 114–127 (2004).

63. R. A. Rodriguez, *et al.*, Photoreactivation of bacteriophages after UV disinfection : Role of genome structure and impacts of UV source. *Water Res.* **55**, 143–149 (2014).

64. R. Sommer, T. Haider, A. Cabaj, E. Heidenreich, M. Kundi, Increased inactivation of Saccharomyces cerevisiae by protraction of UV irradiation. *Appl. Environ. Microbiol.* **62**, 1977 LP – 1983 (1996).

65. K. S. Fallon, T. M. Hargy, E. D. Mackey, H. B. Wright, J. L. Clancy, Development and characterization of nonpathogenic surrogates for UV reactor validation. *J. - AWWA* **99**, 73–82 (2007).

66. S. E. Luria, R. Dulbecco, Genetic Recombinations Leading to Production of Active Bacteriophage from Ultraviolet Inactivated Bacteriophage Particles. *Genetics* **34**, 93–125

(1949).

67. S. E. Beck, H. B. Wright, T. M. Hargy, T. C. Larason, K. G. Linden, Action spectra for validation of pathogen disinfection in medium-pressure ultraviolet (UV) systems. *Water Res.* **70**, 27–37 (2015).

68. D. M. Gunter-Ward, *et al.*, Efficacy of ultraviolet (UV-C) light in reducing foodborne pathogens and model viruses in skim milk. *J. Food Process. Preserv.* **42**, e13485 (2018).

69. Z. Bohrerova, H. Shemer, R. Lantis, C. A. Impellitteri, K. G. Linden, Comparative disinfection efficiency of pulsed and continuous-wave UV irradiation technologies. *Water Res.* **42**, 2975–2982 (2008).

70. M. Otaki, *et al.*, Inactivation differences of microorganisms by low pressure UV and pulsed xenon lamps. *Water Sci. Technol.* **47**, 185–190 (2003).

71. M. R. Templeton, R. C. Andrews, R. Hofmann, Impact of iron particles in groundwater on the UV inactivation of bacteriophages MS2 and T4. *J. Appl. Microbiol.* **101**, 732–741 (2006).

72. M. R. Templeton, R. Hofmann, R. C. Andrews, UV inactivation of humic-coated bacteriophages MS2 and T4 in water. *J. Environ. Eng. Sci.* **5**, 537–543 (2006).

73. E. Timchak, V. Gitis, A combined degradation of dyes and inactivation of viruses by UV and UV/H2O2. *Chem. Eng. J.* **192**, 164–170 (2012).

74. C. Bowker, A. Sain, M. Shatalov, J. Ducoste, Microbial UV fluence-response assessment using a novel UV-LED collimated beam system. *Water Res.* **45**, 2011–2019 (2011).

75. J. J. Cornelis, Z. Z. Su, J. Rommelaere, Direct and indirect effects of ultraviolet light on the mutagenesis of parvovirus H-1 in human cells. *EMBO J.* **1**, 693–699 (1982).

76. D. A. Battigelli, M. D. Sobsey, D. C. Lobe, The Inactivation of Hepatitis a Virus and other

Model Viruses by UV Irradiation. *Water Sci. Technol.* **27**, 339–342 (1993).

77. N. Giese, J. Darby, Sensitivity of microorganisms to different wavelengths of UV light: implications on modeling of medium pressure UV systems. *Water Res.* **34**, 4007–4013 (2000).

78. J. Ho, M. Seidel, R. Niessner, J. Eggers, A. Tiehm, Long amplicon (LA)-qPCR for the discrimination of infectious and noninfectious phix174 bacteriophages after UV inactivation. *Water Res.* **103**, 141–148 (2016).

79. H. Liltved, H. Hektoen, H. Efraimsen, Inactivation of bacterial and viral fish pathogens by ozonation or UV irradiation in water of different salinity. *Aquac. Eng.* **14**, 107–122 (1995).

80. H. Liltved, C. Vogelsang, I. Modahl, B. H. Dannevig, High resistance of fish pathogenic viruses to UV irradiation and ozonated seawater. *Aquac. Eng.* **34**, 72–82 (2006).

81. A. K. Øye, E. Rimstad, Inactivation of infectious salmon anaemia virus, viral haemorrhagic septicaemia virus and infectious pancreatic necrosis virus in water using UVC irradiation. *Dis. Aquat. Organ.* **48**, 1–5 (2001).

82. Y. Ye, P. H. Chang, J. Hartert, K. R. Wigginton, Reactivity of Enveloped Virus Genome, Proteins, and Lipids with Free Chlorine and UV254. *Environ. Sci. Technol.* **52**, 7698–7708 (2018).

83. G. D. Harris, V. D. Adams, D. L. Sorensen, M. S. Curtis, Ultraviolet inactivation of selected bacteria and viruses with photoreactivation of the bacteria. *Water Res.* **21**, 687–692 (1987).

84. J. C. Chang, *et al.*, UV inactivation of pathogenic and indicator microorganisms. *Appl. Environ. Microbiol.* **49**, 1361–1365 (1985).

85. D. Li, A. Z. Gu, M. He, H.-C. Shi, W. Yang, UV inactivation and resistance of rotavirus evaluated by integrated cell culture and real-time RT-PCR assay. *Water Res.* **43**, 3261–3269 (2009).

86. Y. A. Smirnov, S. P. Kapitulets, N. N. Amitna, V. A. Ginevskaya, N. V Kaverin, Effect of UV-Irradiation on rotavirus. *Acta Virol.* **35**, 1–6 (1991).

87. B. R. Wilson, P. F. Roessler, E. Vandellen, M. Abbaszadegan, C. P. Gerba, Coliphage-MS-2 as a UV Water Disinfection Efficacy Test Surrogate for Bacterial and Viral Pathogens in *Water Quality Technology Conference 1992, Parts I and II*, (AMER WATER WORKS ASSOC, 1993), pp. 219–235.

88. P. Huber, B. Petri, S. Allen, J. S. Lumsden, Viral haemorrhagic septicaemia virus IVb inactivation by ultraviolet light, and storage viability at 4 and −20 °C. *J. Fish Dis.* **33**, 377–380 (2010).

89. A. M. de Roda Husman, *et al.*, Calicivirus Inactivation by Nonionizing (253.7-Nanometer-Wavelength [UV]) and Ionizing (Gamma) Radiation. *Appl. Environ. Microbiol.* **70**, 5089 LP – 5093 (2004).

90. B. K. Mayer, H. Ryu, D. Gerrity, M. Abbaszadegan, Development and validation of an integrated cell culture-qRTPCR assay for simultaneous quantification of coxsackieviruses, echoviruses, and polioviruses in disinfection studies. *Water Sci. Technol.* **61**, 375–387 (2010).

91. Z. Zavadova, L. Gresland, M. Rosenbergova, Inactivation of single- and double-stranded ribonucleic acid of enchephalomycarditis virus by ultraviolet light. *Acta Virol.* **12**, 515+ (1968).

92. Q. Zhong, A. Carratala, R. Ossola, V. Bachmann, T. Kohn, Cross-Resistance of UV- or

Chlorine Dioxide-Resistant Echovirus 11 to Other Disinfectants. *Front. Microbiol.* **8** (2017).

93. G. W. Park, K. G. Linden, M. D. Sobsey, Inactivation of murine norovirus, feline calicivirus and echovirus 12 as surrogates for human norovirus (NoV) and coliphage (F+) MS2 by ultraviolet light (254 nm) and the effect of cell association on UV inactivation. *Lett. Appl. Microbiol.* **52**, 162–167 (2011).

94. H. Werbin, R. C. Valentine, A. D. McLaren, Photobiology of RNA Bacteriophages — I. Ultraviolet Inactivation and Photoreactivation Studies. *Photochem. Photobiol.* **6**, 205–213 (1967).

95. T. Tanaka, O. Nogariya, N. Shionoiri, Y. Maeda, A. Arakaki, Integrated molecular analysis of the inactivation of a non-enveloped virus, feline calicivirus, by UV-C radiation. *J. Biosci. Bioeng.* **126**, 63–68 (2018).

96. J. A. Tree, M. R. Adams, D. N. Lees, Disinfection of feline calicivirus (a surrogate for Norovirus) in wastewaters. *J. Appl. Microbiol.* **98**, 155–162 (2005).

97. T. Sigstam, *et al.*, Subtle differences in virus composition affect disinfection kinetics and mechanisms. *Appl. Environ. Microbiol.* **79**, 3455–3467 (2013).

98. L. Guerrero-Latorre, E. Gonzales-Gustavson, A. Hundesa, R. Sommer, G. Rosina, UV disinfection and flocculation-chlorination sachets to reduce hepatitis E virus in drinking water. *Int. J. Hyg. Environ. Health* **219**, 405–411 (2016).

99. J. Lee, K. Zoh, G. Ko, Inactivation and UV Disinfection of Murine Norovirus with TiO(2) under Various Environmental Conditions. *Appl. Environ. Microbiol.* **74**, 2111–2117 (2008).

100. L. F. Batch, C. R. Schulz, K. G. Linden, Evaluating Water Quality Effects on UV

Disinfection of MS2 Coliphage. *J. - AWWA* **96**, 75–87 (2004).

101. B. Sara E., *et al.*, Disinfection Methods for Treating Low TOC, Light Graywater to California Title 22 Water Reuse Standards. *J. Environ. Eng.* **139**, 1137–1145 (2013).

102. S. E. Beck, *et al.*, Comparison of UV-Induced Inactivation and RNA Damage in MS2 Phage across the Germicidal UV Spectrum. *Appl. Environ. Microbiol.* **82**, 1468–1474 (2016).

103. S. E. Beck, *et al.*, Evaluating UV-C LED disinfection performance and investigating potential dual-wavelength synergy. *Water Res.* **109**, 207–216 (2017).

104. Z. Bohrerova, H. Mamane, J. J. Ducoste, K. G. Linden, Comparative Inactivation of Bacillus Subtilis Spores and MS-2 Coliphage in a UV Reactor: Implications for Validation. *J. Environ. Eng.* **132**, 1554–1561 (2006).

105. J. L. Braunstein, F. J. Loge, G. Tchobanoglous, J. L. Darby, Ultraviolet disinfection of filtered activated sludge effluent for reuse applications. *Water Environ. Res.* **68**, 152–161 (1996).

106. E. I. Budowsky, G. V Kostyuk, A. A. Kost, F. A. Savin, Principles of selective inactivation of viral genome. 2. Influence of stirring and optical-density of the layer to be irradiated upon uv-indeuced inactivation of viruses. *Arch. Virol.* **68**, 249–256 (1981).

107. M. A. Butkus, M. P. Labare, J. A. Starke, K. Moon, M. Talbot, Use of aqueous silver to enhance inactivation of coliphage MS-2 by UV disinfection. *Appl. Environ. Microbiol.* **70**, 2848–2853 (2004).

108. M. Cho, V. Gandhi, T.-M. Hwang, S. Lee, J.-H. Kim, Investigating synergism during sequential inactivation of MS-2 phage and Bacillus subtilis spores with UV/H2O2 followed by free chlorine. *Water Res.* **45**, 1063–1070 (2011).

109. H. Guo, J. Hu, Effect of hybrid coagulation–membrane filtration on downstream UV disinfection. *Desalination* **290**, 115–124 (2012).

110. A. H. HAVELAAR, C. C. E. MEULEMANS, W. M. POTHOGEBOOM, J. KOSTER, Inactivation of bacteriophage-MS2 in waste-water effluent with monochromatic and polychromatic ultraviolet-light. *WATER Res.* **24**, 1387–1393 (1990).

111. N. M. Hull, K. G. Linden, Synergy of MS2 disinfection by sequential exposure to tailored UV wavelengths. *Water Res.* **143**, 292–300 (2018).

112. R. M. Jenny, O. D. Simmons, M. Shatalov, J. J. Ducoste, Modeling a continuous flow ultraviolet Light Emitting Diode reactor using computational fluid dynamics. *Chem. Eng. Sci.* **116**, 524–535 (2014).

113. D. Jolis, The Effect of Storage and Lag Time on MS2 Bacteriophage Susceptibility to Ultraviolet Radiation. *Water Environ. Res.* **74**, 516–520 (2002).

114. D. Lénès, *et al.*, Assessment of the removal and inactivation of influenza viruses H5N1 and H1N1 by drinking water treatment. *Water Res.* **44**, 2473–2486 (2010).

115. W. J. Lodder, *et al.*, Reduction of bacteriophage MS2 by filtration and irradiation determined by culture and quantitative real-time RT–PCR. *J. Water Health* **11**, 256–266 (2013).

116. H. Mamane-Gravetz, K. G. Linden, A. Cabaj, R. Sommer, Spectral Sensitivity of Bacillus subtilis Spores and MS2 Coliphage for Validation Testing of Ultraviolet Reactors for Water Disinfection. *Environ. Sci. Technol.* **39**, 7845–7852 (2005).

117. M. J. Mattle, T. Kohn, Inactivation and Tailing during UV254 Disinfection of Viruses: Contributions of Viral Aggregation, Light Shielding within Viral Aggregates, and Recombination. *Environ. Sci. Technol.* **46**, 10022–10030 (2012).

118. E. G. Mbonimpa, E. R. Blatchley III, B. Applegate, W. F. Harper Jr, Ultraviolet A and B wavelength-dependent inactivation of viruses and bacteria in the water. *J. Water Health* **16**, 796–806 (2018).

119. T. J. Nieuwstad, A. H. Havelaar, The kinetics of batch ultraviolet inactivation of bacteriophage-MS2 and microbiological calibration of an ultraviolet pilot plant. *J. Environ. Sci. Heal. PART A-ENVIRONMENTAL Sci. Eng. TOXIC Hazard. Subst. Control* **29**, 1993–2007 (1994).

120. K. A. Sholtes, *et al.*, Comparison of ultraviolet light-emitting diodes and low-pressure mercury-arc lamps for disinfection of water. *Environ. Technol.* **37**, 2183–2188 (2016).

121. D. C. Shoults, N. J. Ashbolt, Total staphylococci as performance surrogate for greywater treatment. *Environ. Sci. Pollut. Res.* **25**, 32894–32900 (2018).

122. M. D. Sobsey, D. A. Battigelli, S. G-A, S. Newland, RT-PCR amplification detects inactivated viruses in water and wastewater. *Water Sci. Technol.* **38**, 91–94 (1998).

123. J. A. Tree, M. R. Adams, D. N. Lees, Virus inactivation during disinfection of wastewater by chlorination and UV irradiation and the efficacy of F+ bacteriophage as a "viral indicator." *Water Sci. Technol.* **35**, 227–232 (1997).

124. Z. Wang, X. Xing, L. Ma, The inactivation effect of ultraviolet disinfection reactor on the high concentration of bacteriophage MS2. *J. FOOD Agric. Environ.* **11**, 1042–1044 (2013).

125. Y. Wang, E. Araud, J. L. Shisler, T. H. Nguyen, B. Yuan, Influence of algal organic matter on MS2 bacteriophage inactivation by ultraviolet irradiation at 220 nm and 254 nm. *Chemosphere* **214**, 195–202 (2019).

126. S. Weng, *et al.*, Infectivity reduction efficacy of UV irradiation and peracetic acid-UV

combined treatment on MS2 bacteriophage and murine norovirus in secondary wastewater effluent. *J. Environ. Manage.* **221**, 1–9 (2018).

127. K. R. Wigginton, B. M. Pecson, T. Sigstam, F. Bosshard, T. Kohn, Virus Inactivation Mechanisms: Impact of Disinfectants on Virus Function and Structural Integrity. *Environ. Sci. Technol.* **46**, 12069–12078 (2012).

128. S. Rattanakul, K. Oguma, Inactivation kinetics and efficiencies of UV-LEDs against Pseudomonas aeruginosa, Legionella pneumophila, and surrogate microorganisms. *Water Res.* **130**, 31–37 (2018).

129. Y. A. Smirnov, S. P. Kapitulez, N. V Kaverin, Effects of UV-irradiation upon Venezuelan equine encephalomyelitis virus. *Virus Res.* **22**, 151–158 (1992).

### Supplementary Figures

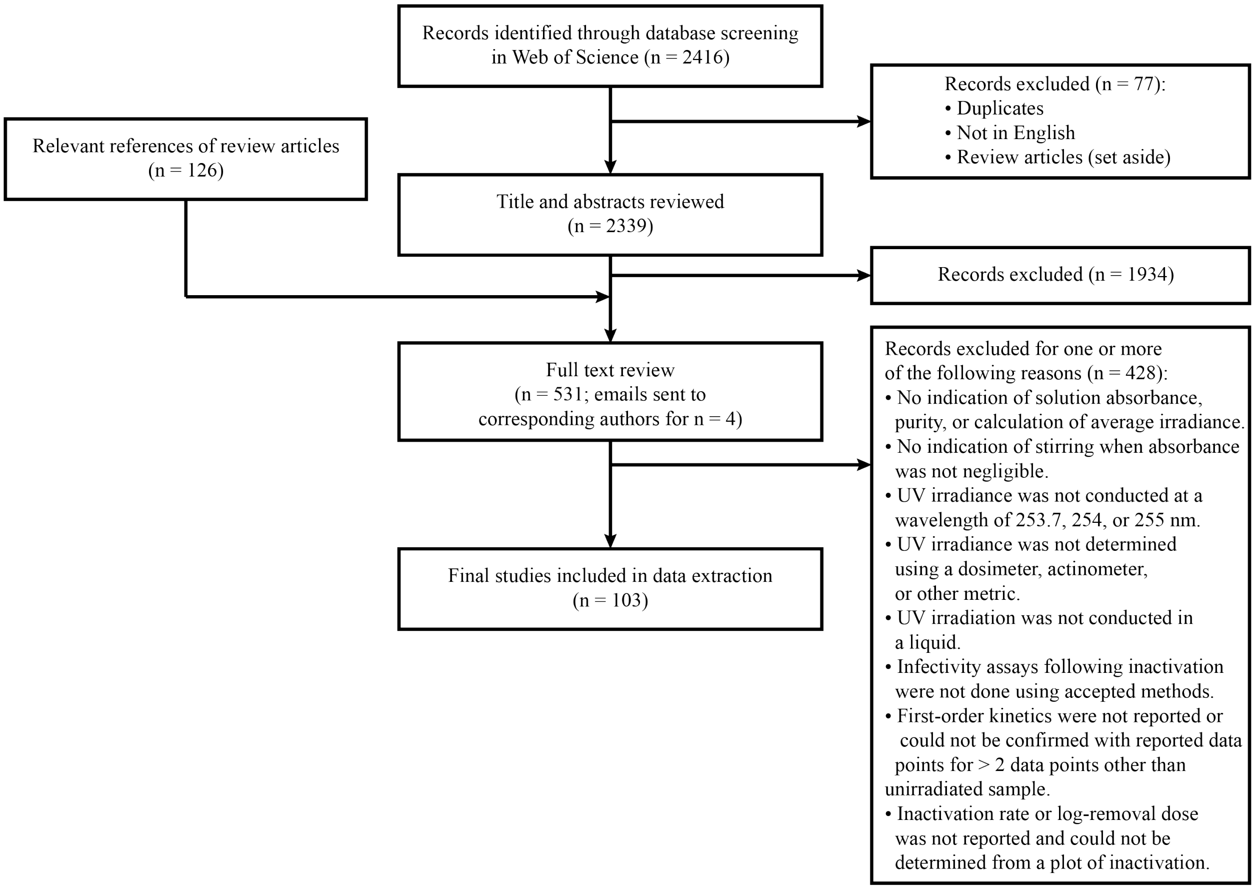


***Figure S1***. *Flow chart of rapid systematic literature review conducted to collect high-quality $UV_{254}$ virus inactivation rate constants.*

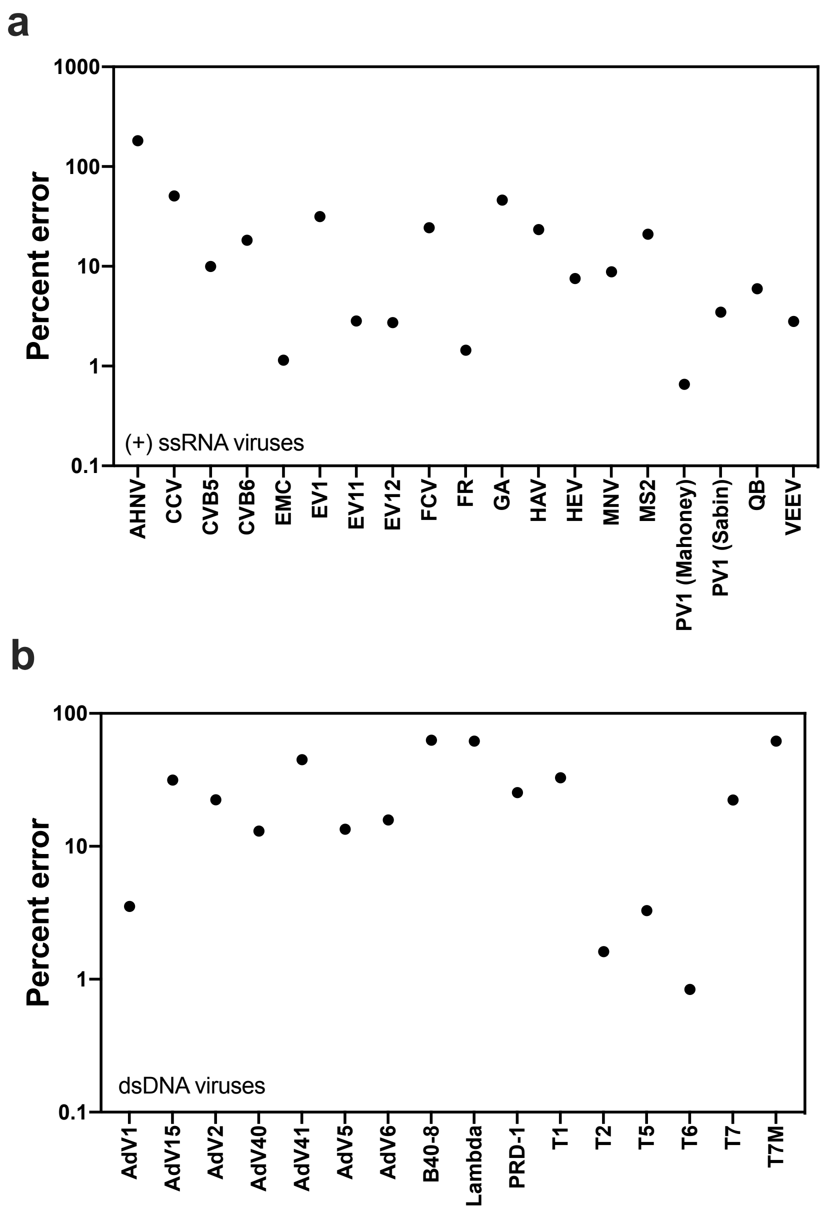


***Figure S2.*** *Percent error of the predicted inactivation rate constant from the mean experimental rate constant for each virus where the predicted constants were determined using the top performing (+) ssRNA virus model (a) and dsDNA virus model (b).*

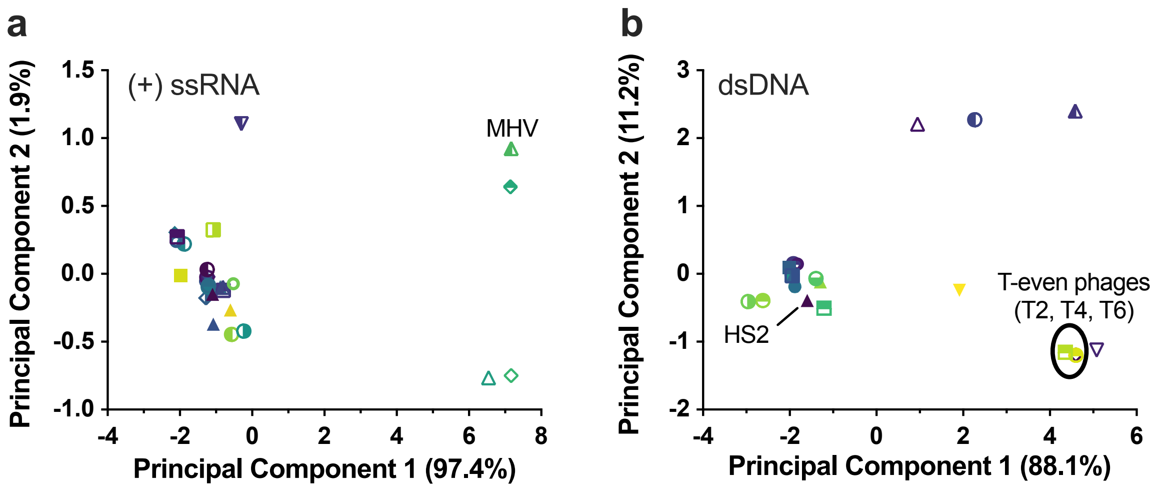


***Figure S3.*** *Principal component analyses of virus genome attributes for (+) ssRNA viruses (a) and dsDNA viruses (b).. Viruses included all viruses from the systematic review with full genome sequence information, and viruses used in predictions. Principal component analyses were conducted on standardized genome attributes, as described in methods.*

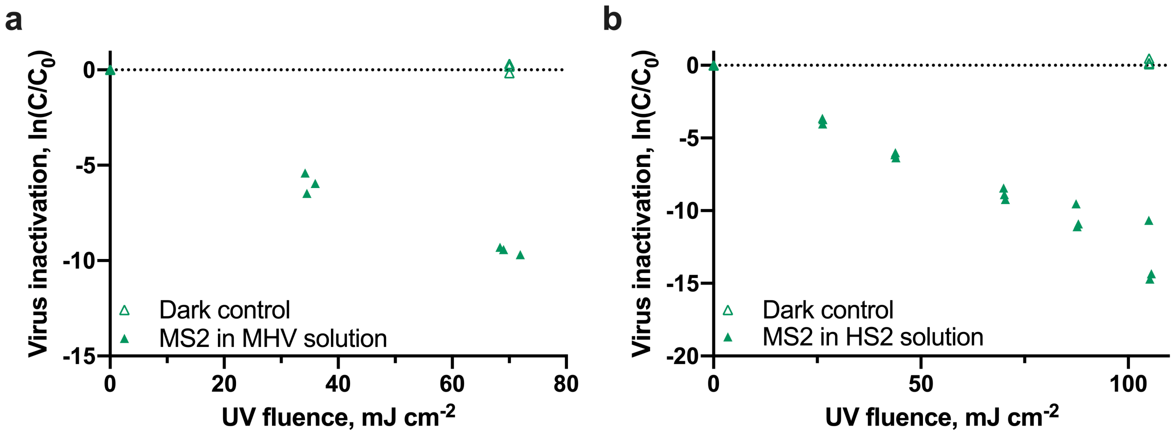


***Figure S4.*** *Inactivation of MS2 following $UV_{254}$ irradiation when MS2 was in the MHV experimental solution (a) and the HS2 experimental solution (b). Independent replicates (N = 3) are shown for each fluence. MS2 was spiked into the experimental solution at a final concentration of $10^9$ pfu/mL.*

#### Supplementary Tables

**Table S1. Experimental $UV_{254}$ inactivation rate constants extracted from the rapid systematic literature review and virus rate constants and weights used in modeling work.**
*This table is provided as an external data file.*

**Table S2. Virus genome sequence sources and predictor information for all viruses used in training/validation and prediction.**
*This table is provided as an external data file.*

**Table S3. Model performance metrics for top-performing models of each class for each subset of viruses used in the training and validation set.**
*This table is provided as an external data file.*

**Table S4. Results of pairwise multiple linear regression model comparisons.**
*This table is provided as an external data file.*

**Table S5. Results of pairwise model comparisons.**
*This table is provided as an external data file.*

**Table S6. Predicted virus inactivation rate constants from the top performing dsDNA virus model and top performing (+) ssRNA virus model.**
*This table is provided as an external data file.*
